# Structural basis for saccharide binding by human RNase 2/EDN, a protein combining enzymatic and lectin properties

**DOI:** 10.64898/2026.03.20.713198

**Authors:** Xincheng Kang, Guillem Prats-Ejarque, Ester Boix, Jiarui Li

## Abstract

Human RNase 2 (eosinophil-derived neurotoxin, EDN) is a major eosinophil granule protein of the vertebrate-specific RNase A superfamily and is involved in antiviral response and inflammation. Identifying ligand-binding pockets in EDN is thus relevant to structure-based drug design. In our laboratory we identified by protein crystallography a conserved site at the protein surface binding to carboxylic anion molecules (malonate, tartrate and citrate). Searching for potential biomolecules rich in anion groups and considering previous report of EDN binding to glycosaminoglycans, we explored the protein binding to saccharides. Next, EDN crystals were soaked with mono- and disaccharides, and the 3D structures of ten complexes were solved by X-ray crystallography at atomic resolution. We identified protein binding pockets to glucose, fucose, mannose, sucrose, galactose, trehalose, N-acetyl-D-glucosamine, N-acetylmuramic acid, and the sialic acid N-acetylneuraminic acid. A main site for glucose, fucose, and galactose was located adjacent to the spotted carboxylic anion site. Secondarily, N-acetylneuraminic acid, N-acetylmuramic acid, sucrose, galactose, and mannose shared another protein surface region. Overall, the saccharides clustered into seven defined sites, outlining a conserved recognition pattern, which was further analysed by molecular modelling. Interestingly, within the RNase A family, we find amphibian RNases that were initially isolated as carbohydrate binding proteins and named as leczymes, combining enzymatic and lectin properties. The present data is the first systematic structural characterization of a mammalian sugar-binding RNase within the family. The results highlight unique EDN residues that mediate its sugar specific interactions, of particular interest for a better understanding of the protein physiological role.

**Highlights:**

- structure of RNase 2 in complex with mono and disaccharides at atomic resolution
- identification of RNase 2 unique sugar binding sites
- characterization of a mammalian RNase A family enzyme with lectin properties

**Graphical Abstract:** 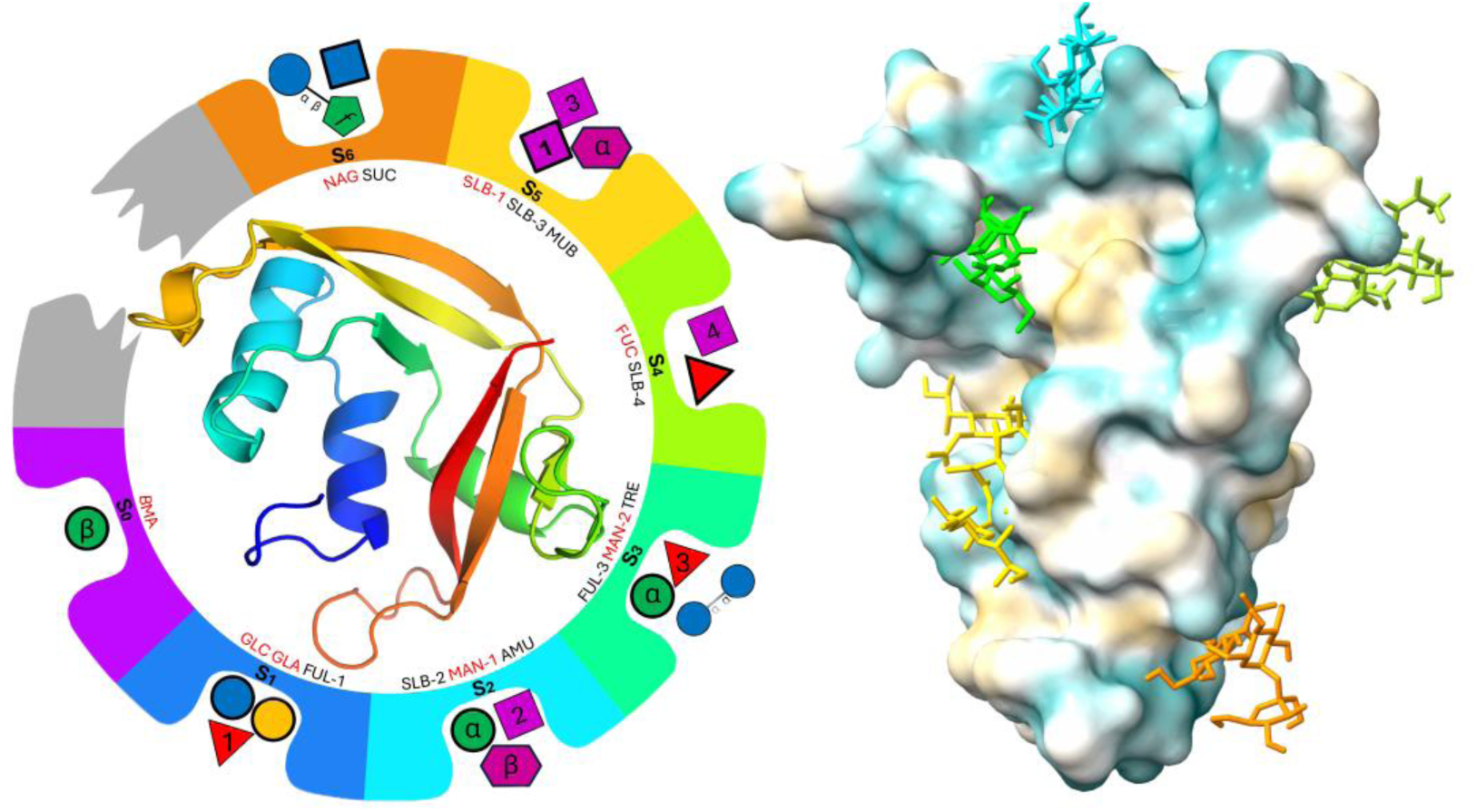

## 1. Introduction

Human RNase 2 belongs to the RNase A superfamily, which includes 8 canonical types. The RNase A family members participate in host defence and can be defined as multifaceted proteins [1-3]; in addition to their catalytic ribonucleolytic action, a variety of biological properties have been reported [4]. RNase 2, or the Eosinophil-Derived Neurotoxin (EDN), is one of the main proteins stored in the eosinophil secretory granules and takes its name upon its discovery from its neurotoxicity when injected to guinea pigs [5]. On the other hand, several amphibian ribonucleases were initially isolated as carbohydrate-binding proteins during the 70s and named as lectins [6, 7]. Upon the discovery of the first frog sialic acid-binding lectin (cSBL), the authors proposed the term leczyme, in reference to its dual nature: sugar binding and RNase catalytic activity. The protein was isolated from the oocytes of *Rana catesbeiana* and its lectin properties were associated to its antitumoral action. Curiously, the frog cSBL is not homologous to any other family of lectin proteins [7]. Other homologous frog RNases were characterized with selective agglutinating activities for tumour cells [8]. Nitta et al showed that the frog RNase antitumor activity was mediated by interaction with sialoglycoproteins exposed at cancer cell membranes [9]. The authors also observed that the protein internalization was followed by inhibition of cell proliferation. However, the SBLs lectin-glycan binding proteins still remain poorly characterized. On its side, RNase 2 is a mammalian member of the RNase A family member of particular interest, as it originated, together with its close homologue RNase 3, from a duplication event that took place during primate evolution, after divergence of New Word from Old World monkeys [10]. RNase 3 (also called Eosinophil Cationic Protein; ECP), shares a 70% identity with RNase 2. Both RNases are abundant in eosinophils and stored within their secretory granules[11], with distinct antimicrobial properties and expression patterns. To note, eosinophil RNase 3/ECP is associated to airway inflammation and has been routinely used to monitor asthma and other hyperinflammatory diseases [4]. On its side, EDN expression is triggered by viral infections [12] and has also been associated to exacerbated response to infection [13], or even to rare cases of blood disorders linked to RNA vaccination [14]. Accordingly, EDN was proposed as a biomarker for some hyperinflammation conditions and blood disorder pathologies[15].

The eosinophil RNases branch, which has undergone an unusual rapid evolutionary pace within primates, is of particular interest to study human innate immunity and has also attracted special attention for structure-based guided protein design [16]. Interestingly, both eosinophil RNases were reported to display affinity for heparin and interact mostly through electrostatic interactions between arginine surface exposed residues and the glycosaminoglycan (GAG) acid groups [17-19]. Later research demonstrated how addition of heparin could prevent EDN interaction with bronchial epithelial Beas-2B cells, suggesting that binding to some exposed saccharide molecules at the cell surface could mediate the protein action [20]. By protein-cell interaction studies using oligosaccharides of increasing length, the authors identified an heparin heptamer as the most efficient inhibitor of the protein binding [20]. Next, by site-directed mutagenesis and in silico predictions, the GAG interaction was mostly attributed to protein cationic residues and sulphate groups [21]. Likewise, ECP was also reported to present high affinity to GAGs and work by site directed mutagenesis, NMR and molecular modelling allowed the identification of protein residues involved in binding to heparin and heparan sulphate derivates [18, 19, 22-24].

Recently, a renewed interest on RNases potential binding to sugar derivates aroused with the unexpected discovery of glycoRNAs [25]. We can speculate that the presence of glycoRNAs exposed at blood cells surface and participating in cell-cell recognition [26] might mediate some of the eosinophil protein biological roles. Thus, it is of great interest to understand the protein binding mode to cell surfaces and specifically to saccharide units.

In the present work we describe EDN complexes with a wide variety of monosaccharides derivates, together with two disaccharides and three carboxylate anions, all solved by X-ray crystallography (MX) at atomic resolution. Results illustrate unambiguously the protein specific binding mode to each sugar type. MX combined with molecular modelling has enabled us to outline a multisite architecture at the protein surface conformed by specific hydrophobic patches, together with polar interactions, unique for EDN-saccharide recognition.

## 2. Methodology

### 2.1. Abbreviations

We list below the used sugar names and corresponding abbreviations. Table 1 lists the common name, specific name, and the corresponding three-letter code according to the Protein Data Bank (PDB) nomenclature.

**Table 1.**
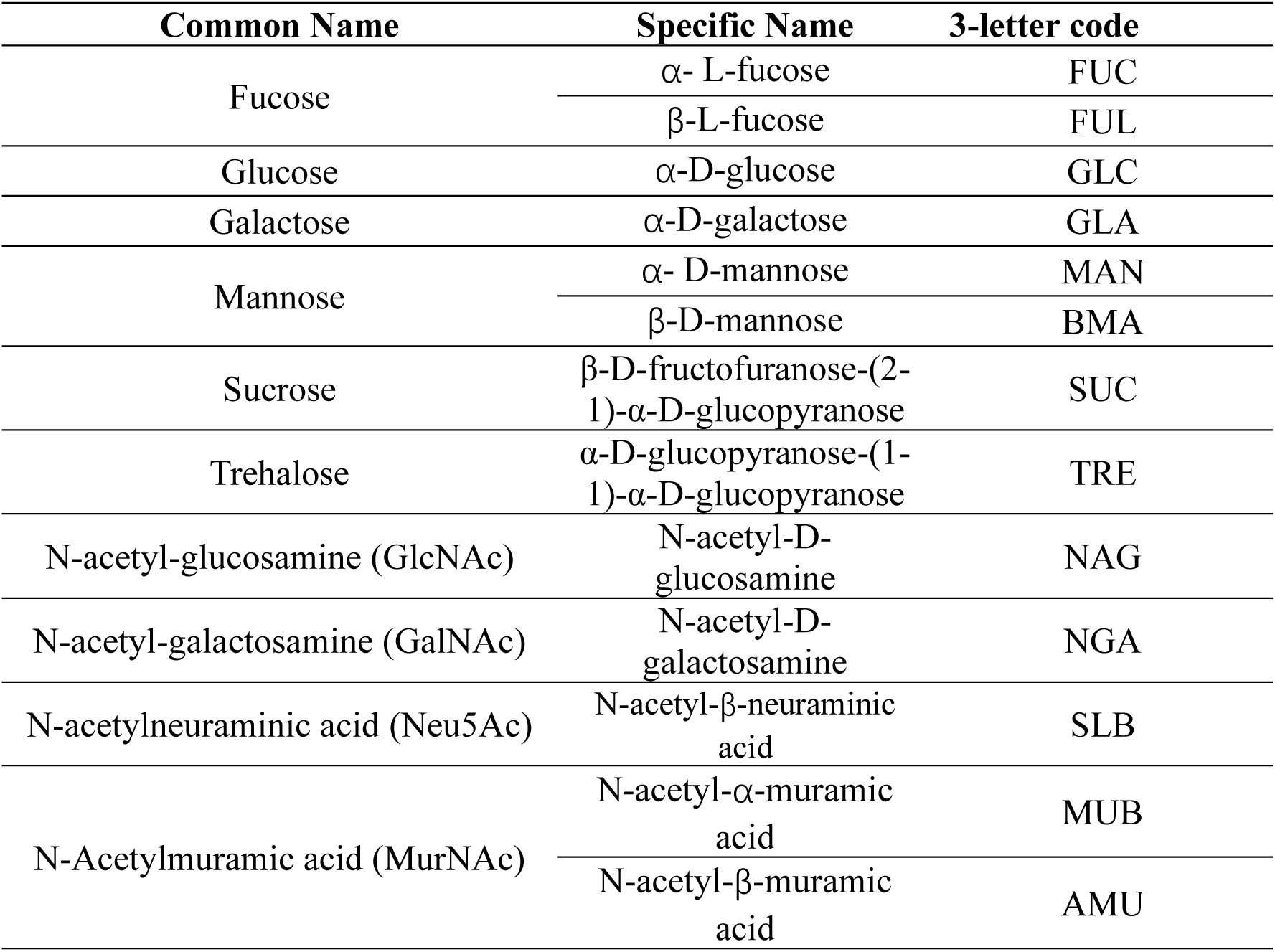
List of used abbreviations for saccharides.

### 2.2 Materials

Carbohydrates used for crystal soaking comprised: D(+)-sucrose (CAS 57-50-1; ITW Reagents), D-trehalose dihydrate (CAS 6138-23-4; Apollo Scientific); D-(+)-glucose (CAS 50-99-7; Merck), D-(+)-galactose (CAS 59-23-4; Glentham), L-(-)-fucose (CAS 2438-80-4; Thermo Fisher), D-(+)-mannose (CAS 3458-28-4; Apollo Scientific); N-acetyl-D-glucosamine (CAS 7512-17-6; TCI), N-acetyl-D-galactosamine (CAS 14215-68-0; Glentham), N-acetylneuraminic acid (CAS 131-48-6; Apollo Scientific), and N-acetylmuramic acid (CAS 10597-89-4; Bachem).

### 2.3 Protein expression and purification

Recombinant RNase 2 was expressed and purified following the previously established protocol in our laboratory [27]. Briefly, *E. coli* BL21(DE3) cells were transformed with the pET11c vector including the EDN gene for protein expression. Following IPTG addition, cells were harvested, lysed and processed to isolate inclusion bodies. Solubilization of inclusion bodies was achieved using 100 mM Tris-acetic acid, 2 mM EDTA, and 6 M guanidine hydrochloride (pH 8.5). Next, 80 mM of reduced glutathione was added, and samples were incubated under a nitrogen atmosphere for two hours at room temperature. Protein refolding was initiated by a 100-fold dilution into refolding buffer (0.1 M Tris-HCl, 0.5 M L-arginine·HCl, 0.2 mM oxidized glutathione, pH 8.5) and maintained for 72 hours at 4° C. The buffer was then exchanged to 150 mM sodium acetate (pH 5). Final purification was conducted on an FPLC system using a Resource S cation exchange column. The protein was eluted with a linear gradient of NaCl (0–2 M) in 150 mM sodium acetate buffer (pH 5). Purified fractions were concentrated, desalted and lyophilized for long-term storage at -20° C, and assessed for purity using 15% SDS-PAGE.

### 2.4 Crystallization conditions

Lyophilized EDN was dissolved in a buffer containing 10 mM Tris-HCl, pH 7.4 and 50 mM NaCl and adjusted to a final concentration of 5 mg/mL and subjected to initial crystallization screening using the sitting-drop vapour-diffusion method. For each condition, 100 nL of protein solution was mixed with 100 nL of reservoir solution and the plates were incubated at 18 °C. Initial screening were performed using commercial kits, including Index HR2-134, JCSG-plus HT-96 (MD1-40/40-FX), PACT premier HT-96 Eco Screen (MD1-36-ECO), PEG HR2-139, and Structure Screen 1+2 HT-96 (MD1-30), encompassing a total of 480 distinct conditions. Crystals were observed in 106 wells. Crystallization hits were predominantly associated with carboxylate-containing salts, including formate, acetate, malonate, citrate, tartrate, succinate, and malate, frequently combined with PEG 3350 (16–25% w/v) as precipitant. Successful conditions typically spanned pH 6.0–8.5 and involved buffers such as HEPES, Tris, or Bis-Tris propane. Conditions were selected from the initial screening hits and refined, as described below.

Conditions that yielded crystals were systematically refined by varying the pH, salt and precipitant concentrations to obtain single, high-quality crystals and establish mother-liquor conditions compatible with ligand soaking. Three crystallization hits that produced abundant crystals and had a good resolution (1.20Å -1.30Å) were selected for optimization (see Table S1). Based on condition 1, which yielded the highest number of crystals after three months incubation at 18℃, we pursued subsequent optimization. Last, we selected the following crystallization condition: 0.22 M sodium formate, 0.1 M Bis-Tris propane (pH 8.7), and 12% (w/v) PEG 3350, which yielded high-quality crystals within 1–3 weeks by incubation at 18°C.

### 2.5 Sugar soaking conditions

Crystals of suitable size were selected for ligand-soaking experiments. For standard soaking conditions, a solution was prepared by mixing 0.6 µL of reservoir solution with 0.4 µL of a 50%(w/v) sugar stock. Crystals were transferred to this solution and incubated for a defined period (see Table S2), then harvested and flash-cooled in liquid nitrogen.

For saccharide derivatives, such as N-acetyl-D-glucosamine (NAG), 5-N-acetyl-β-neuraminic acid (SLB), N-Acetyl-D-galactosamine (NGA) and N-acetylmuramic acid (MUB), the soaking protocol was adapted to overcome solubility limitations and the relatively high salt concentration of some crystallization buffers. In these cases, the monosaccharide derivatives were dissolved directly in 25% PEG 3350 and 0.1M Bis-Tris propane, followed by heating at 37°C for 10 minutes and centrifuging for 10 minutes to collect the supernatant, which was used as the soaking solution. EDN crystals were soaked in 1 µL of this solution for up to 4 days.

### 2.6 X-ray data collection, data processing and structure refinement

Diffraction data were collected at the XALOC BL13 beamline station of ALBA synchrotron using a λ = 0.9795 Å. Data collection was performed at 100K using a Pilatus3 X 6M detector 100Hz (Dectris®, Switzerland), images were taken at t_exp_ = 0.1s, Δφ = 0.1°, and the total oscillation range per dataset was 180°. All crystals diffracted at atomic resolution (0.94 to 1.3 Å). The raw data were processed using the automated autoPROC pipeline[28]. Within this pipeline, indexing and integration were performed by XDS[29], while data scaling and merging were carried out with AIMLESS[30] using the *CCP4* suite[31]. Refinement was performed with *Phenix* [32]using the ligand-free protein structure (PDB ID 1GQV[33]) as the starting model for molecular replacement, subsequent model building and manual adjustments were carried out in Coot[34]. Iterative cycles of refinement and rebuilding were applied until convergence, at which point water molecules and ligands were included on the basis of unambiguous difference map electron density. Data processing and structure solving statistics for all structures is summarized in Table S2.

### 2.7 Molecular modelling

Molecular docking was conducted using *AutoDock Vina-Carb v1.0* [35] to evaluate carbohydrate binding at the sites identified in the crystal structures and to predict whether the characterized protein binding-site architecture could accommodate an extended oligosaccharide.

Protein receptor coordinates were taken from the corresponding crystal structures. Prior to docking, crystallographic water molecules were removed, and the receptor was prepared using *AutoDockTools(ADT)v1.5.6*[36] by adding hydrogen atoms and assigning partial compute Gasteiger charges. For docking calculations, the prepared receptor was converted to PDBQT format.

All monosaccharide ligands were downloaded from *PubChem*[37]. The oligosaccharide (polymeric) ligand model was generated using the CSDB *GlycanBuilder* and GLYCAM[38, 39]. *ADT* was used to convert ligands to the PDBQT format and to set the identified active torsions, allowing for full conformational flexibility during the docking process.

Docking was conducted in a semi-flexible manner, with the receptor treated as rigid and the ligands as fully flexible. LigPlot+ Interaction diagrams [40] were applied to identify residues surrounding each crystallographically observed sugar-binding site. The docking search space (grid box) was centred on these residues but deliberately expanded beyond the minimal interacting envelope to avoid bias from overly restrictive sampling. Box dimensions were adjusted according to ligand size and typically extended by approximately 25% beyond the residue boundaries to ensure sufficient conformational freedom while maintaining focus on the experimentally validated binding pocket (see Table S3). Docking boxes and corresponding configuration files were generated using *ADT*.

Docking calculations were performed using *AutoDock Vina-Carb*. The receptor was treated as rigid, whereas ligands were modelled with full torsional flexibility as defined in *ADT*. The exhaustiveness parameter was adjusted according to ligand size. For monosaccharide docking, the exhaustiveness parameter was set to 300. To balance sampling thoroughness with computational efficiency as molecular complexity increased, this value was adjusted to 150 for di- and trisaccharide docking and further reduced to 50 for the heptasaccharide. This tiered approach accounts for the significantly expanded conformational space of larger glycans. Each docking run generated nine binding poses (num modes = 9), and the energy range was set to 5 kcal/mol. To ensure reproducibility and sampling consistency, each ligand was docked independently using three different random seeds. Binding poses were ranked according to predicted binding energy. For the heptasaccharide, the final pose was selected by comparison with the carbohydrate-binding sites previously identified by X-ray crystallography. Among the nine Vina-Carb poses, the selected model reproduced the largest number of shared hydrogen-bonding and van der Waals (vdW) interactions with the experimentally observed bound sugars at this site, while also showing a favourable binding energy. The selected pose was extracted and combined with the receptor coordinates in *PyMOL* to generate the protein-glycan complex, and the glycan topology was verified against the original heptasaccharide model.

## 3. Results

We obtained crystals of human RNase 2/EDN in complex with several mono- and disaccharides: α-glucose (GLC), α/β-fucose (FUC/FUL), α-galactose (GLA), α/β-mannose (MAN/BMA), sucrose (SUC), trehalose (TRE), N-acetyl-D-glucosamine (NAG), N-acetylmuramic acid (MUB), and 5-N-acetyl-β-neuraminic acid (SLB); together with protein complexes with carboxylate anions: malonate (MLI), tartrate (TLA) and citrate (CIT). All datasets were collected at atomic resolution; data-processing, refinement and structure validation statistics are summarized in Table S2. Coordinates and structure factors have been deposited at the PDB under the accession codes: 8QEW, 9GNT, 9GO7, 9GO8, 9GO4, 9GQ0, 9GQM, 9GQP, 9QYO, 9QR7, 9R6D, 9RAN and 9R6E. Overall, two anion-binding sites (A1, A2) and seven sugar-binding sites (S_0_-S_6_) were identified at the protein surface.

### 3.1 A novel anion binding site

When searching for new crystallization conditions for EDN, we obtained high quality crystals for diverse ones. Interestingly, when solving the protein 3D structures in some conditions we identified bound anion salts. Thus, we obtained the protein in complex with a variety of mono, di and tricarboxylate anions up to six carbons, i.e.: acetate, malonate, tartrate and citrate. All complexes were solved at atomic resolution showing clear unambiguous electron density for ligand positioning (Figure S1). Results highlight that most of the ligand molecules clustered into a unique site, showing overlapping of acetate, malonate, tartrate and citrate molecules (Figure 1). Curiously, despite acetate was only present in the crystallization buffer from 9GQM dataset, an acetate ligand bound at the same location was identified in other datasets (PDB ID: 9GO4, 9GO8, 9GQ0, 9GQM, 9RAN, 9R6D, 9GQP, 9QR7, and 9R6E). Considering that acetate was absent from the crystallization buffer used in these conditions, the results suggests that acetate would come from the buffer used during the protein purification procedure. Surprisingly, the bound acetate was not removed even after sample desalting and buffer exchange against water before lyophilization step. Interestingly, the acetate molecule overlapped onto one of the carboxyl groups of citrate, malonate and tartrate. Overall, interactions at this main anion binding site (named as A1) are provided by residues T46, N50, V54, G75, S76, V78 and Y107 (Figure 1). We observe conserved hydrogen bonds between T46 and Y107 side chain OH donor groups to carboxylate O acceptors. Besides, the cavity is formed by hydrophobic contacts with side chains of N50, V54 and V78, and additional hydrogen bonds between NH main chain of G75 and S76 with carboxylate groups.

**Figure 1.**
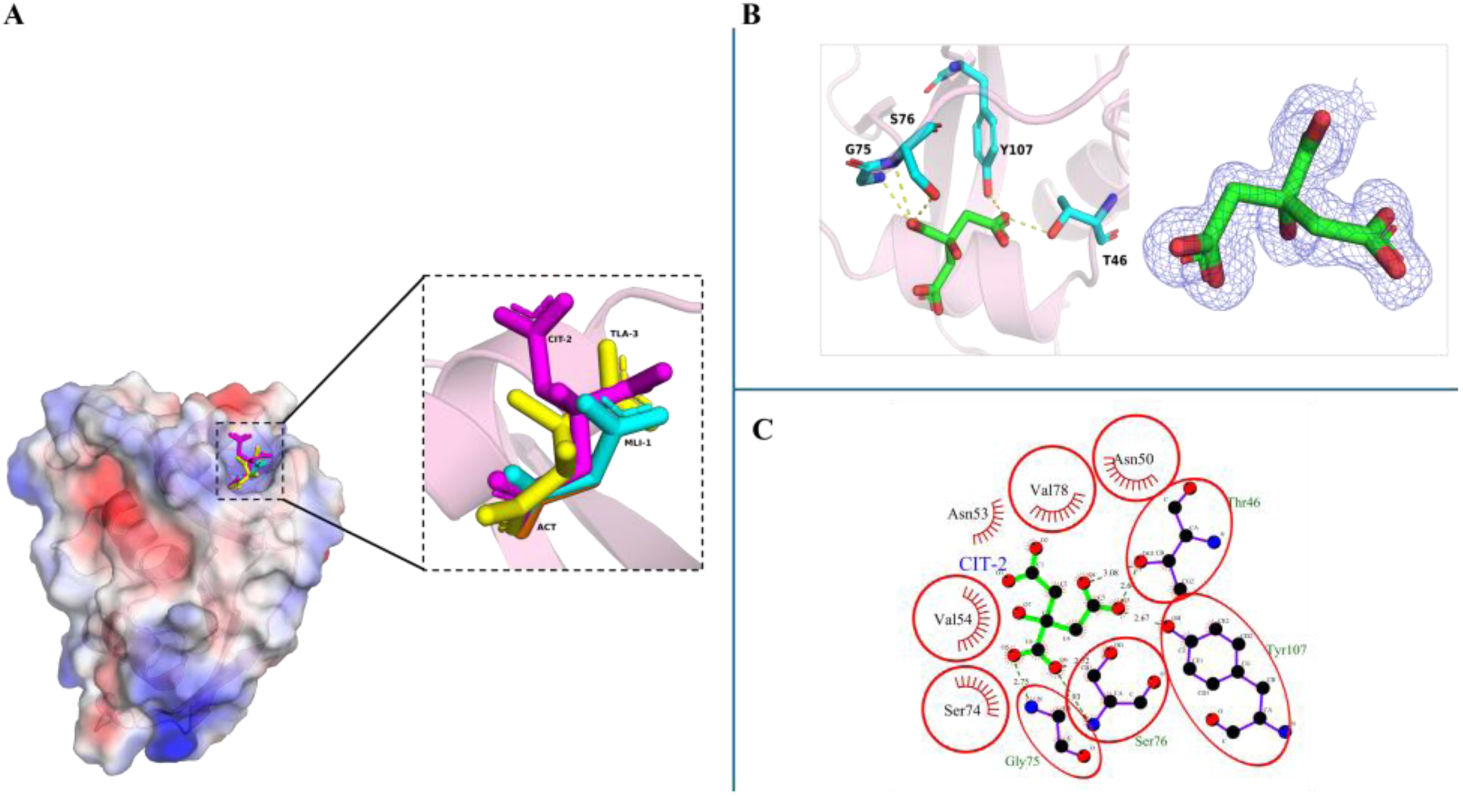
Structural analysis of EDN complexes with acetate (ACT), tartrate (TLA), citrate (CIT), and malonate (MLI). (A) Overall view showing the clustering of the 4 anions at the A1 site. (B) Close-up of the CIT binding pocket with its 2FoFc electron density map. Interacting residues are displayed as sticks, and H-bonds are indicated by yellow dashed lines. (C) Ligplot interaction diagram for CIT. Green dashed lines denote hydrogen bonds. Residues highlighted by red ellipses (e.g., T46, S76 and Y107) are consistently observed across different protein-ligand complexes, identifying them as common binding hotspots.

Overall, we can see conservation of hydrogen bonds to 2 carboxylate groups, where T46 and Y107 share direct interactions with the same carboxylate group. Secondarily, we found a second cluster at the active site binding pocket, conformed by residues Q14, H15, K38, H129 and L130, showing a second molecule of acetate, tartrate and citrate, which we named A2 (Figure S2).

### 3.2 Novel saccharide binding sites for EDN

Next, based on the observation of the EDN conserved binding mode to the carboxylate groups, we decided to explore the biological significance of the spotted surface binding region. Thus, we considered the potential protein interaction with biological molecules offering anion groups for interaction. Considering that EDN was reported to have high affinity binding to glycosaminoglycans[20] and some RNase A family members were characterized as lectins[6, 7], we decided to soak EDN crystals with a variety of saccharide compounds.

We firstly screened the protein potential binding to neutral sugars including glucose, fucose, galactose, mannose, sucrose, trehalose by soaking EDN crystals. Overall, we successfully obtained structures at atomic resolution of EDN binding to: one glucose (α-D-glucopyranose, GLC; PDB ID: 9GNT), four fucoses (one α-L-fucopyranose, FUC and three β-L-fucopyranose, FUL-1-3; PDB ID: 9GQ0), one galactose (α-D-galactopyranose, GLA; PDB ID: 9RAN), three mannoses (two α-D-mannopyranoses, MAN-1/MAN-2 and one β-D-mannopyranose, BMA; PDB ID: 9QYO), one sucrose (β-D-fructofuranose-(2-1)-α-D-glucopyranose, SUC; PDB ID: 9GO7) and one trehalose (α-D-glucopyranose-(1-1)-α-D-glucopyranose, TRE; PDB ID: 9R6D) (Figure S3).

Interestingly, we observed that the main binding region for glucose, glucose and galactose partly overlaps with the first previously identified carboxylic anion site (Figure 2). This specific site for sugars, adjacent to the A1, was named as S_1_. For both glucose and galactose, the O6 hydroxyl group hydrogen bonds to the backbone carbonyl oxygen of Q77, whereas the O1 hydroxyl group interacts with the carboxyl group of the adjacent anion salt. We observed the presence of a bound citrate from the crystallization condition, in the case of GLC (Figure 2B and C) and an acetate in case of GLA complex (Figures 3 and S4). Besides, in GLA, O1 also mediates an additional hydrogen bond with the -NH_2_ group of the N50 side chain. When fucose binds to EDN as a ligand, the side chain hydroxyl group of T46 forms a hydrogen bond with the fucose O3 group (Figure S4). For all three ligands, residues L45, V78, and P79 help to stabilize the binding pocket through hydrophobic interactions.

**Figure 2.**
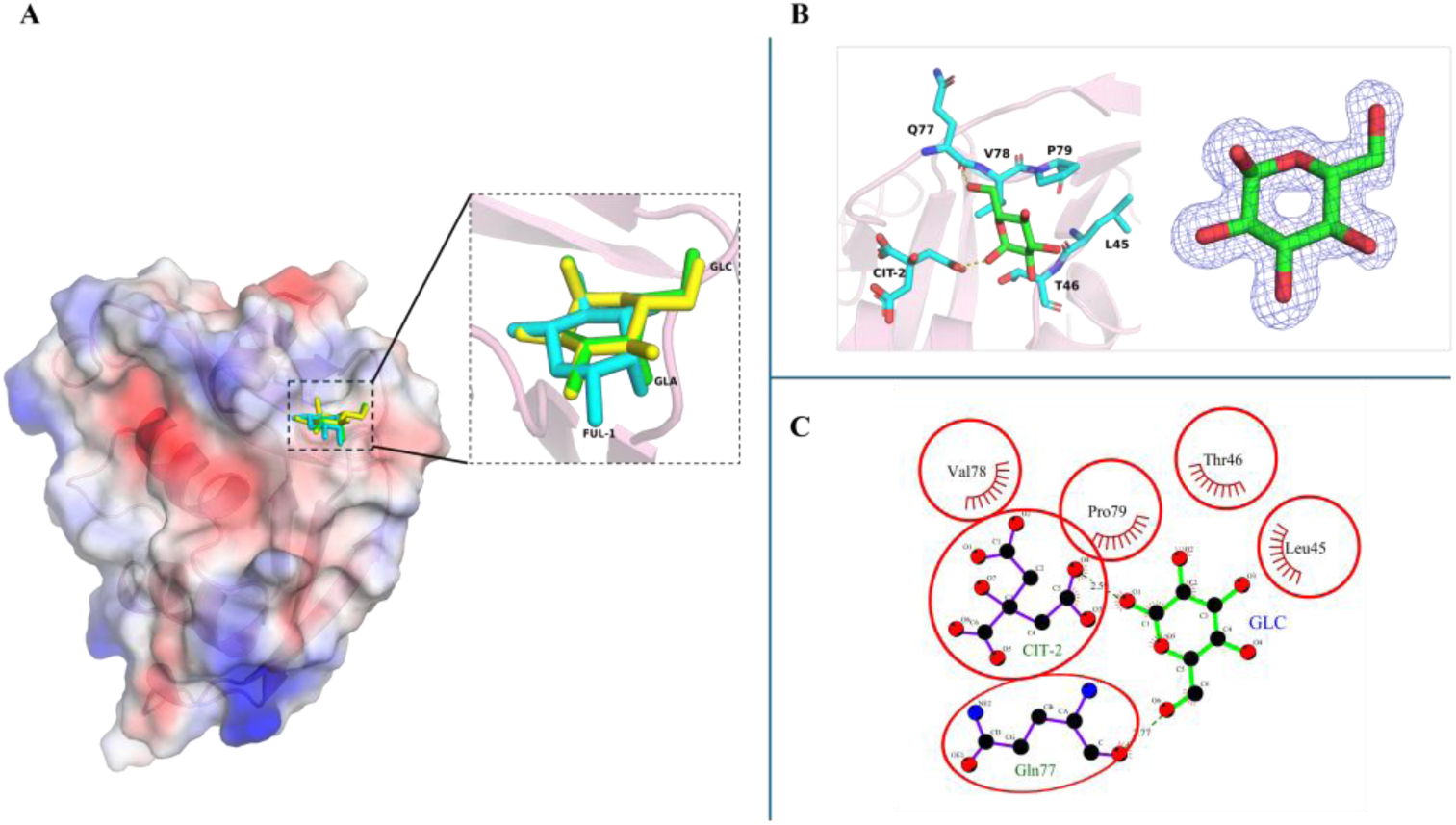
Structural analysis of EDN complexes with glucose (GLC), fucose (FUL-1), and galactose (GLA). (A) Overall view showing the clustering of the three sugars at the S_1_ site. (B) Close-up of the GLC binding pocket with its electron density map. Interacting residues are displayed as sticks, and hydrogen bonds are indicated by yellow dashed lines. (C) Ligplot interaction diagram for GLC. Green dashed lines denote hydrogen bonds. Residues highlighted by red ellipses are consistently observed across different ligand complexes, identifying them as common binding hotspots.

**Figure 3.**
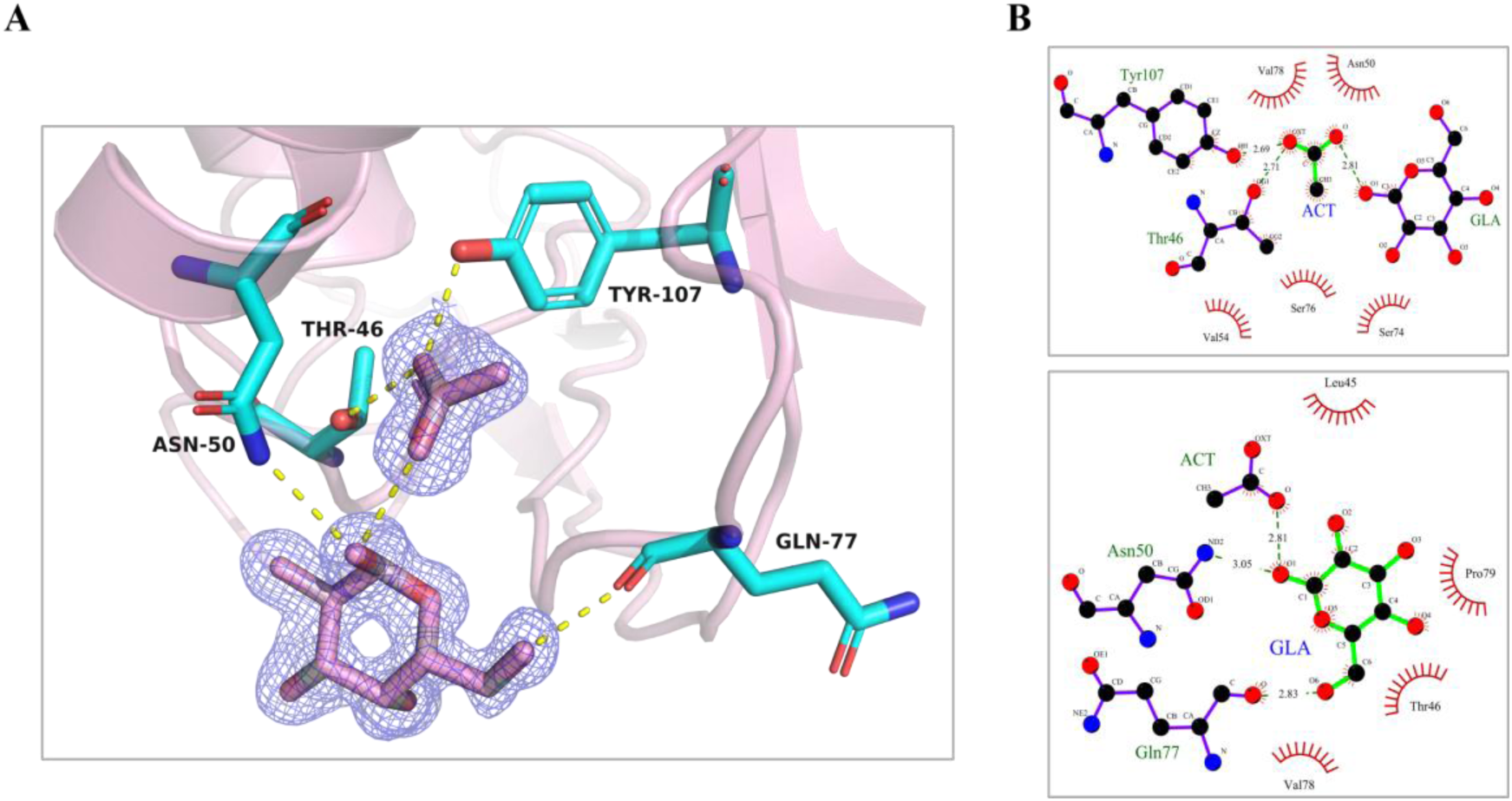
Location of galactose (GLA) at the S_1_ site and acetate (ACT) at A1 site in EDN complex. (A) 2FoFc electron density map for both ligands and protein residues involved in hydrogen bond interactions. (B) The interactions between the ligand and the surrounding amino acid residues are shown by LigPlot diagrams, dashed lines representing hydrogen bonds and red arcs representing hydrophobic interactions.

Further analysis of solved structures also revealed the presence of other binding sites. We identified another site for β-mannose (BMA) at the left site of the main site, which we named S_0_ (Figure S5A). This site is primarily composed of residues L45, P79, I81, and P102, which provide vdW contacts. In addition, another binding site for α-mannose (MAN-1) was identified, which forms hydrogen bond interactions with H72 and N59, this site was named S_2_ (Figure S6A).

Next, another site was identified, conformed by overlapping of MAN-2, FUL-3 and TRE, named as S_3_ (Figure 4). This site is mainly conformed by residues R68 and N70, that provide H-bonds, together with hydrophobic binding by A110 and H129. This site partially overlaps with the secondary base site B_2_, that accommodates purines in EDN and all other studied RNase A family members [41].

**Figure 4.**
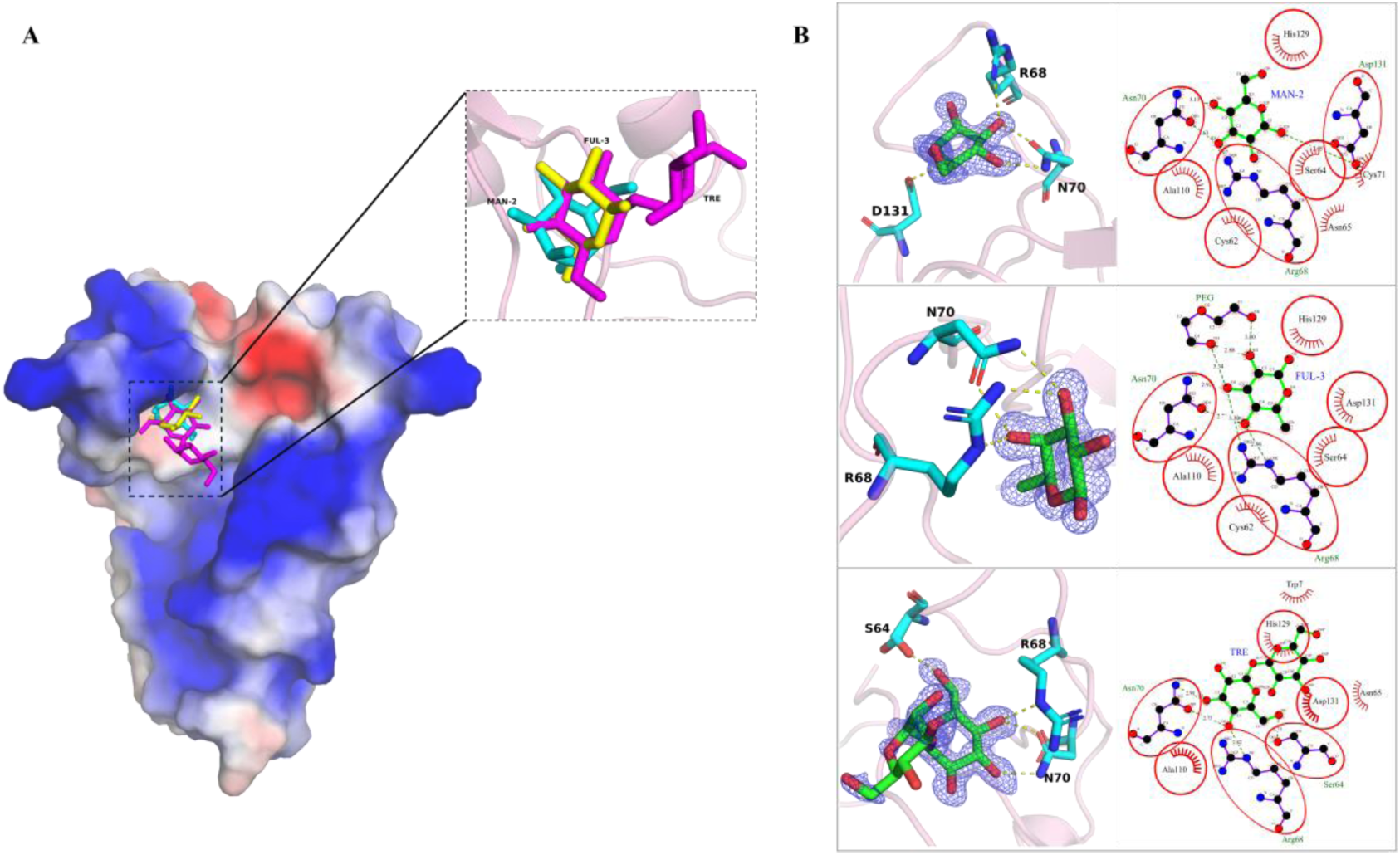
Structural analysis of EDN complexes with mannose (MAN-2), fucose (FUL-3), and trehalose (TRE). (A) Overall view showing the clustering of the three sugars at the S_3_ site. (B) Close-up view of sugar binding at the S_3_ site and LigPlot interaction diagrams. Interacting residues are displayed as sticks; residues highlighted by red ellipses are consistently observed across different ligand complexes, identifying them as common binding hotspots.

Inspection of trehalose complex indicated that for the disaccharide only one of the two monomer units was visible at the electron density difference map, where the GLC unit overlapped with the monosaccharides located at S_3_ site (Figure 4).

Moreover, a complex obtained by soaking crystals simultaneously with both combined ligands confirmed the binding selectivity for each sugar type. Thus, EDN crystals soaking with GLC+ FUC compounds (PDB ID: 9GQM) confirmed GLC binding to S_1_ and FUC to S_4_ site.

Following, based on the observation of closeness of S_1_ and A1, we decided to explore the potential binding of saccharide derivates that incorporate some carboxyl substituents at the pyranose ring. Therefore, we prepared some soaking conditions for the following compounds: N-acetyl-D-glucosamine, the sialic acid N-acetylneuraminic acid, N-acetyl-D-galactosamine and N-acetylmuramic acid. We obtained structures at atomic resolution of EDN binding to one N-acetyl-D-glucosamine (NAG; PDB ID: 9R6E), four N-Acetylneuraminic acids (N-acetyl-β-neuraminic acid, SLB-1/SLB-2/SLB-3/SLB-4; PDB ID: 9GQP) and two N-acetylmuramic acids (one N-acetyl-α-muramic acid, MUB and one N-acetyl-β-muramic acid, AMU; PDB ID: 9QR7) (Figure S7).

First, we identified the AMU and SLB-2 molecules overlapping with the previous site for MAN-1 (S_2_), where the ND2 atom of N59 and the NE2 atom of H72 exhibited shared interactions in all the three sugar molecules (Figure S6). Besides, we observed hydrogen bonds between the OD1 of the D112 side chain and the O4 atom of SLB-2 or the N2 atom of AMU, respectively. Additionally, there was a hydrogen bond interaction between the N5 atom of SLB2 and the OD1 atom of N113. Apart, the binding pocket was further stabilized through hydrophobic contacts mediated by N70 and C111. On the other hand, we found another N-acetylneuraminic acid molecule (SLB-4) at the S_4_ site, overlapping with previously located FUC molecule. In addition, SLB-1, SLB-3, and MUB share a distinct binding pocket, designated as S_5_ (Figure 5A). Last, the NAG molecule overlaps with the previously identified SUC site, sharing the S_6_ binding pocket (Figure 5B).

**Figure 5.**
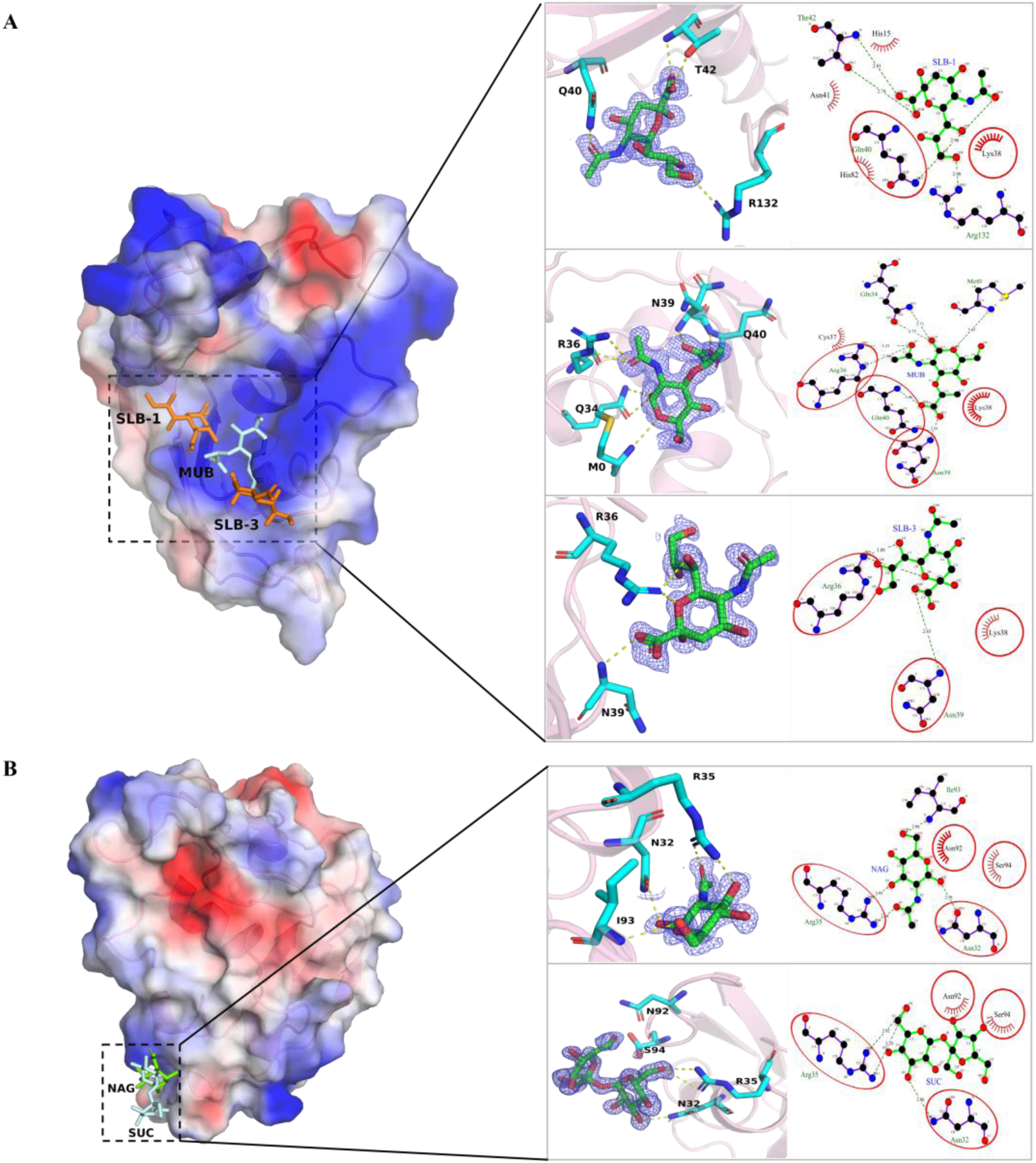
EDN Sugar-binding sites S_5_ and S_6_. (A) Location of the S_5_ site in EDN and interaction diagrams showing the binding of three ligands (SLB-1, SLB-3, and MUB) at the protein surface and their interactions with surrounding amino acid residues. (B) Location of the S_6_ site in EDN and interaction diagrams showing the binding of NAG and SUC with protein residues.

In summary, four bound molecules of SLB were identified at the protein surface, where SLB-1 bound at S_5_ was the one providing a higher number of H-bond and vdW interactions (Figure 5A). A second molecule (SLB-2) also provided abundant specific interactions (Figure S6), whereas much fewer interactions were observed for SLB-3 (Figure 5A). Last, SLB-4 overlaps with FUC at S_4_ (Figure S5B).

On the other hand, soaking with N-acetyl α/β muramic (MUB/AMU) allowed us to confirm the selectivity of N-acetyl saccharides for S_2_ and S_5_, with overlapping with binding sites for the two main SLB molecules. MUB/AMU shares some common features with NAG and SLB, such as the N-acetyl group, but has unique characteristics, such as the lactic C3 chain, with a CH3 group. However, no direct protein interactions with the muramic CH3 group were identified in EDN complexes. Considering muramic unique biological source from bacteria, in contrast to N-acetyl neuraminic acid, most abundant in mammals, the present data does not suggest any specific structural determinants that would mediate a selective protein recognition of bacterial envelope. On the other hand, the N-acetyl group from MUB, shared by NAG and SLB is not recognized by EDN by equivalent interactions. Although AMU shares the same S_2_ cavity with SLB-2, showing in both molecules the recognition of COOH group by complementary interactions by N59 and H72, both compounds have a distinct orientation, with no common features. For example, the N atom of the N-acetyl group is recognized in SLB2 by N113, versus H-bond to D112 in AMU. On its side, the N atom from N-acetyl group from NAG is not directly fixed by any protein residue. The other bound MUB molecules shares S_5_ cavity with SLB-1, but again there is no equivalent interactions, with COOH groups from each molecule recognized differently. Also, in both MUB and SLB-1 we do not identify direct contacts with the N atom from N-acetyl group. However, we do observe specific interactions of R132 with the OH of the C6 glycol group for SLB-1. Another interesting feature of the S_5_ site is its partial overlapping with the main base site B_1_, specific for pyrimidine binding in all RNase A family members, with the participation of T42 that provides a double anchoring point. Complementarily, at the S_5_ sugar site, we also observe the participation of R36, N39 and Q40, residues previously reported to be key for phosphate anchoring at the p_-1_ site[33, 41].

In summary, taking together all the identified sites from the solved protein complexes with the tested mono and disaccharides, we can outline 7 protein regions for sugar binding (S_0_ to S_6_), where S_1_, close to A1, is chosen as the main reference site (Figure 6). A list of all the protein interacting residues involved in each sugar binding site is included in Table 2.

**Figure 6.**
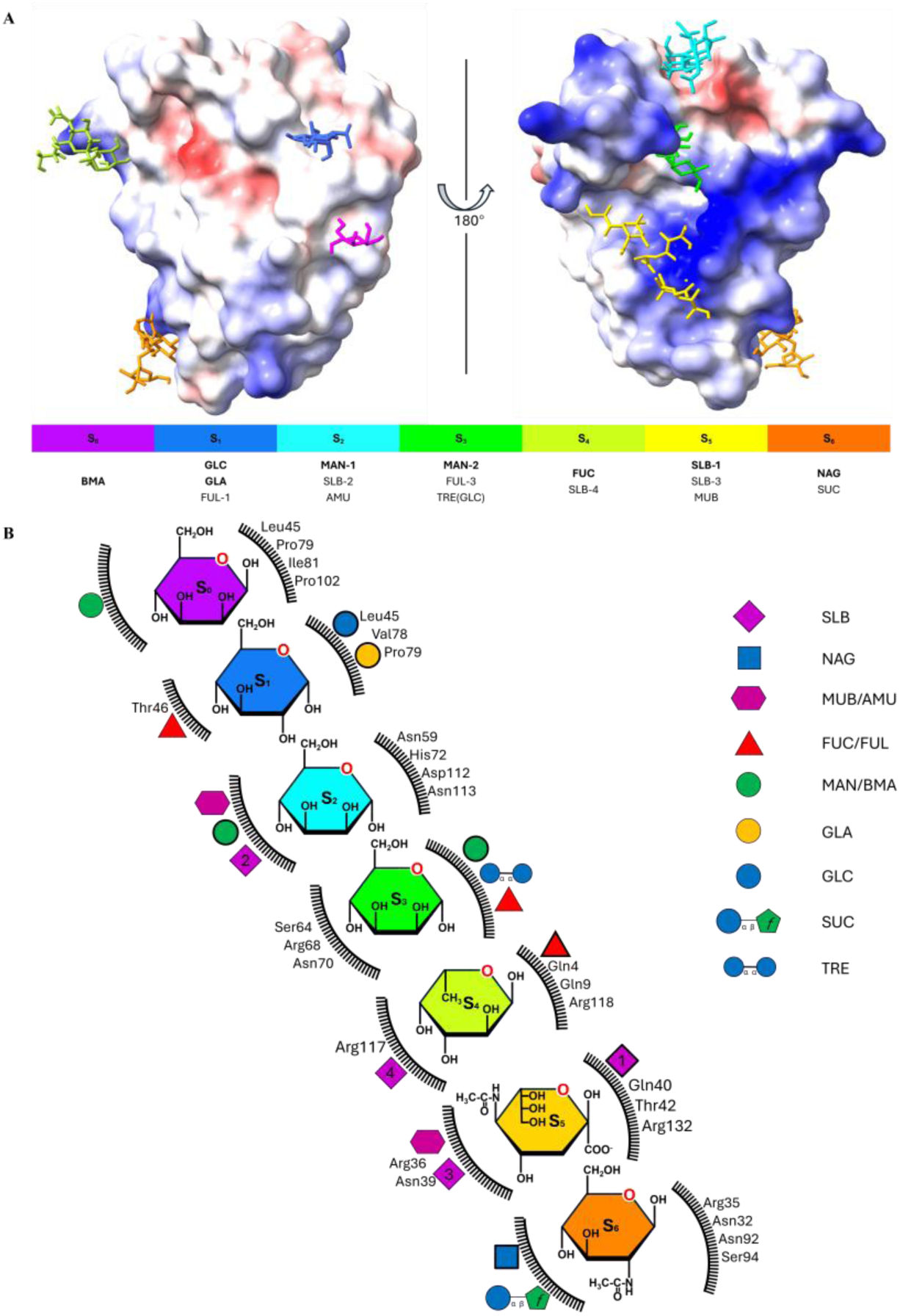
Overview of EDN sugar binding sites. (A) Summary of the locations of S_0_-S_6_ sites at EDN surface structure. The colour of each ligand is consistent with the colour bar below the panel. Abbreviations for the sugars bound at each site are positioned under the colour bars, where bold type highlights the major binding saccharide. (B) Diagram of the identified sugar-binding sites in EDN. The centre shows the mainly sugar bound at each site, with its colour corresponding to panel A. All bound sugars are represented by saccharide symbols located outside the serrated arcs, following the SNFG (Symbol Nomenclature for Glycans) convention. The main interacting amino acid residues for each site are indicated below the respective symbols.

**Table 2.**
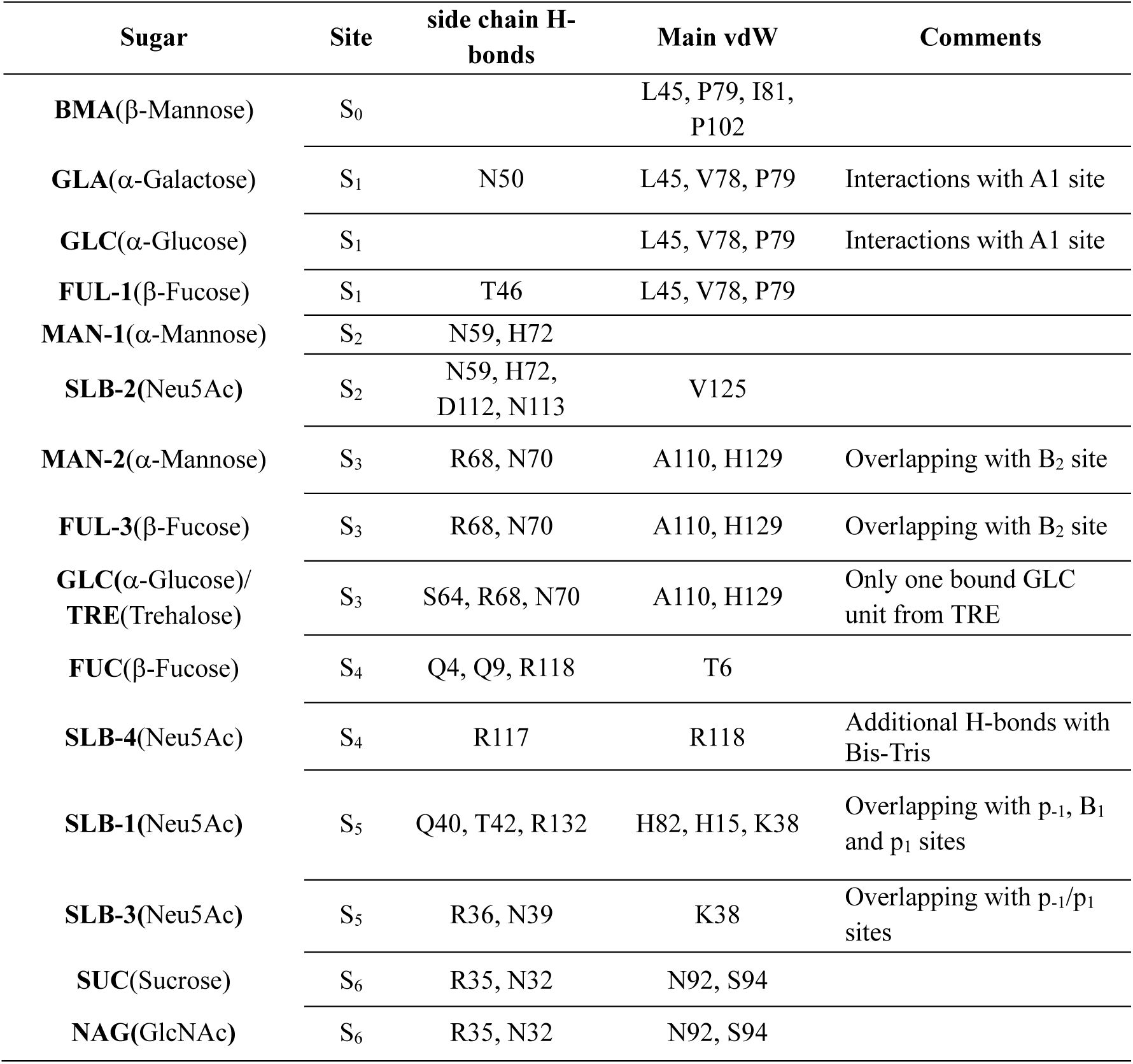
Summary of EDN sugar-binding sites and protein–ligand interactions.

Based on an overall analysis of each binding site, considering the number and specificity of interactions, we designed for each site a principal ascribed sugar (Table 3). Accordingly, we can ascribe GLC/GLA as the main sugars for S_1_. Next, we can locate three mannose binding sites at S_0_, S_2_ and S_3_ respectively. Then, from the 4 fucose molecules bound to EDN, we highlight the one occupying the S_4_ site, observed in both the fucose-only and Fucose+Glucose co-soaking experiments. Last, in case of S_5_ we only observe binding of sialic acids and in S_6_ we find best accommodation for the NAG molecule.

**Table 3.**
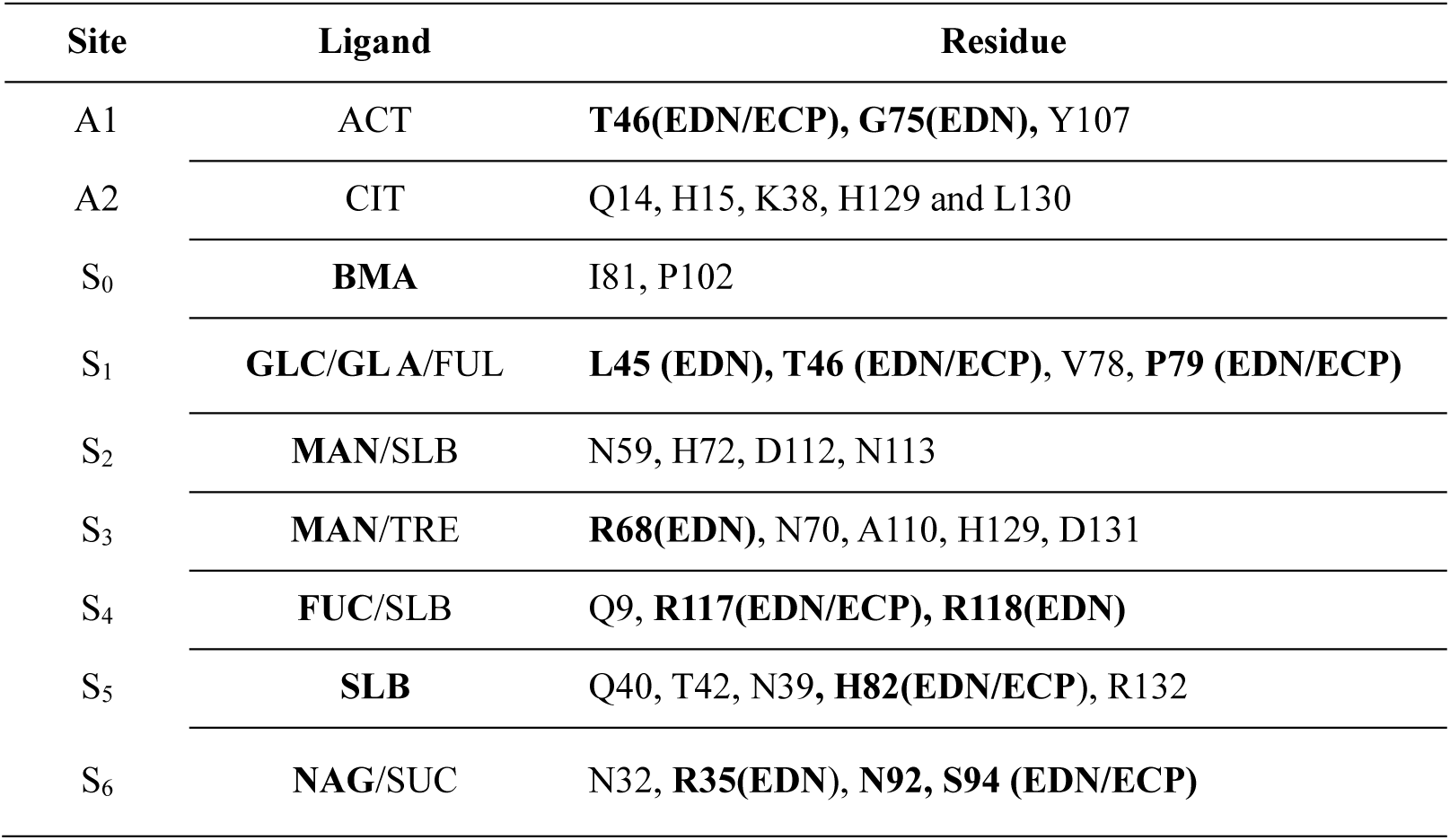
Representative ligands and defining residues for EDN anion- and sugar-binding sites. Main binding ligand and residues unique to EDN or shared with its close homologue ECP are indicated in bold.

Finally, we evaluated the singularity of the identified anchoring sites in EDN respect to their human RNase type counterparts (Figure 7A). We also explored the conservation scale of the spotted residues by use of the *Consurf* server[42] (Figure 7B), together with the *Structure Motif Search* tool from the RCSB Protein Data Bank server.

**Figure 7.**
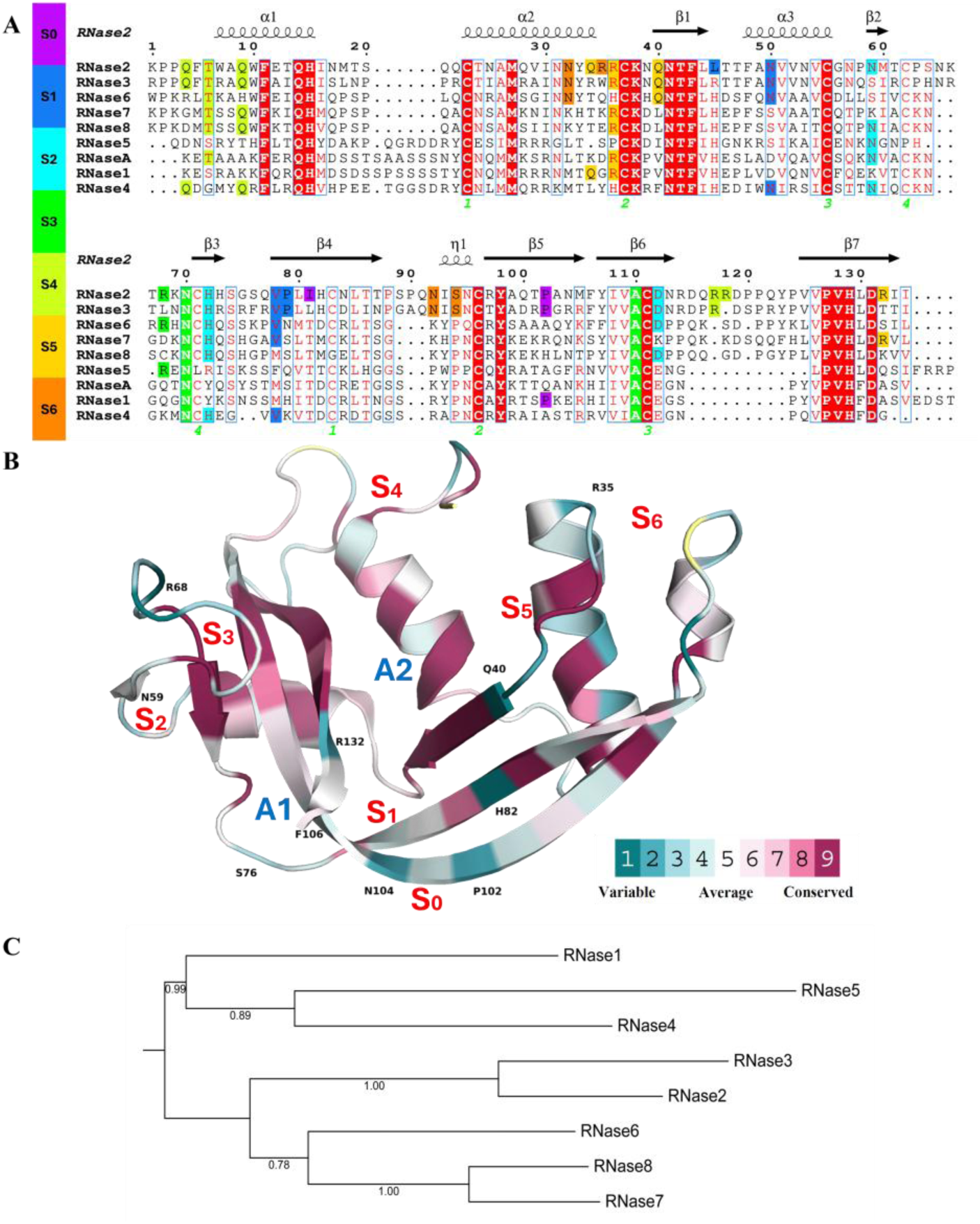
Evolutionary analysis of EDN sugar-binding pockets within the RNase A superfamily. (A) Multiple sequence alignment of the eight canonical human RNases (RNases 1 to 8), with RNase 2/EDN shown on top. Residues contributing to the sugar-binding sites (S_0_–S_6_), as defined from our crystal structures, are indicated by coloured side bars on the left. The secondary-structure elements of EDN are drawn on top of the sequence alignment. (B) Ribbon representation of the EDN structure showing the location of the seven sugar-binding pockets (S_0_–S_6_, red labels) and the two anion sites (A1–A2, blue labels). Residue colouring to highlight evolutionary conservation among the eight human RNases, as obtained from *Consurf* analysis[42], highlighting positions that are representative to EDN. (C) Phylogenetic tree of the eight human RNase A family members. The phylogenetic tree was constructed using the Neighbour-Joining method in MEGA12.

Based on these criteria, we classified each site as: i) unique to EDN (S_1_, S_6_ and A1); ii) shared by the two eosinophil RNases EDN and ECP (S_4_-S_5_) or iii) shared with other family members (S_0_, S_2_, S_3_ and A2) (Table 3). Interestingly, some sites partially overlap with the RNase A family characterized substrate binding site. The S_3_ and A2 site corresponds to the RNase active site, which will anchor the phosphate at p_1_ corresponding to the substrate cleavable phosphodiester bond. We then find overlapping at B_1_ and B_2_ base sites, which include some specific residues for both eosinophil RNases, together with unique ones to EDN. Accordingly, we can ascribe B_1_ to S_5_, B_2_ to S_3_, and p_-1_ associated to S_5_ (see Table 2). Overall, we observe that some sites, such as S_0_, S_1_, S_4_ and S_6_, have unique features to EDN, whereas S_5_ is shared by both EDN and ECP and S_2_ and S_3_ could be found in other family member types (Table 3). Mostly, the sites shared with other RNases are partially overlapping with nucleotide binding sites, such as base binding sites B_1_, B_2_ and phosphate binding p_-1_ site. To summarize, based on the crystallographic data, we defined 7 sugar-binding sites for EDN (see Figures 6 and 7).

### 3.3 Modelling studies to explore EDN binding to oligosaccharides

Next, we applied molecular modelling with the final aim to outline EDN binding site architecture to extended oligosaccharides. Based on the S_0_ to S_6_ sites identified by X-ray crystallography (Figure 6), we decided to explore the protein putative interactions to an heptasaccharide ligand. Beforehand, we performed semi-flexible docking for each monosaccharide ligand at the corresponding EDN binding site identified in the crystal structures. This approach was useful to assess whether the simulations could reproduce the experimentally observed binding modes. All calculations were carried out with *AutoDock Vina-Carb*, with the docking box encompassing all relevant residues highlighted in the LigPlot diagrams. Overall, docking results largely reproduced the experimentally observed binding sites. All estimated binding energies for monosaccharides fall within a similar value range (Table S4). Overall, most sugars show relatively favourable binding energies, particularly at pockets S_2_, S_3_, S_5_ and S_6_, where predicted binding energies are below −4 kcal/mol. In contrast, ligands bound at S_0_, S_1_ and S_4_ generally exhibit slightly weaker interactions, with energies above −4 kcal/mol. Notably, pockets S_2_, S_3_, S_5_ and S_6_ are clustered at the same face of the protein, indicating a higher density of potential carbohydrate-binding sites on that side. For sucrose, crystallographic data indicated that the glucose (GLC) moiety binds more tightly than the fructosyl residue, as reflected by the full refined occupancy (∼1.0) of the GLC ring compared with the lower occupancy (∼0.59) of the fructose moiety and poor electron density. The docking results are consistent with this observation and reproduce the preferential binding of the GLC unit. Four FUC binding sites were also identified in crystal structures, and the corresponding docking scores show slight variations among sites. Altogether, most sugars converge to positions consistent with the crystallographic binding pockets. Overall, we find a good correlation except for the fucose ligand corresponding to FUL-2 location, which failed to reproduce the experimentally identified binding region and, in fact, was not assigned to any of the seven defined pockets by MX. For FUC, GLA, and SLB-3/SLB-4, only part of the docking result locations corresponded to the crystallographic sites, whereas the remaining sugars consistently reproduced the experimentally observed binding positions. At the S_1_ site, although GLA shows a slightly stronger predicted binding energy than GLC, all GLC docking poses converge to the crystallographically observed position, while only a subset of GLA poses correspond to that site.

To further investigate how these monosaccharide sites might contribute to the recognition of extended ligands, we attempted to construct an oligosaccharide model. Based on the macromolecular crystallography (MX) results, a total of 7 binding pockets were defined (Figure 6), encompassing a total of 17 ligands bound to EDN. Some of these binding pockets can accommodate three different ligands (S_1_, S_2_, S_3_ and S_5_), whereas S_0_ contains a single ligand. Within each binding pocket, we selected the dominant sugar monomer based on the number of interactions with protein residues and subsequently constructed a heptasaccharide model using the CSDB *GlycanBuilder* and GLYCAM. We first selected oligosaccharide fragments consisting of monosaccharides from adjacent sugar binding sites, then performed molecular docking at the corresponding sugar binding sites. The results showed that disaccharide fragments exhibited slightly better binding affinity than monosaccharides, specifically the best estimated affinities were: -3.5 kcal/mol for MAN-GLC (S_0_-S_1_), -6.0 kcal/mol for MAN-SLB (S_3_-S_5_), -2.1 kcal/mol for MAN-MAN-SLB (S_2_-S_3_-S_5_), -5.3 kcal/mol for GLA-SLB (S_1_-S_2_), -4.7 kcal/mol for GLA-NAG (S_6_), and -0.6 kcal/mol for NAG-GLA-SLB (S_5_-S_6_). As the sugar chain length increased, the binding affinity gradually increased. Next, we combined these monosaccharides into a heptasaccharide for docking, which yielded 9 solutions, with 4 models clustering together, mainly located within the S_2_, S_3_, S_5_ site region, with individual affinity values below 4 kcal/mol. However, this was not an ideal result, possibly because the spacing between the 7 sugar binding pockets on the EDN surface was significantly larger than the typical spacing between two adjacent sugar residues, making it technically challenging to construct a single oligosaccharide capable of spanning all S_0_–S_6_ sites. Therefore, the docked fragments could not simultaneously reproduce all monosaccharide positions observed in the crystals and failed to fully simulate a potential target with biological significance. In contrast, shorter chains could better simulate sugar binding positions, as evidenced by the good clustering of all models in the aforementioned disaccharide docking results, showing conserved hydrogen bonds with key protein residues identified by MX. To better evaluate the docking results, we selected the three representative regions that could optimize the coverage of all sugar binding sites identified by MX: the heptasaccharide mainly clustered at S_2_, S_3_, S_5_ (Figure 8), MAN-GLC clustered at S_0_ and S_1_ (Figure S8), and NAG-GLA-SLB clustered at S_5_ and S_6_ (Figure S9).

**Figure 8.**
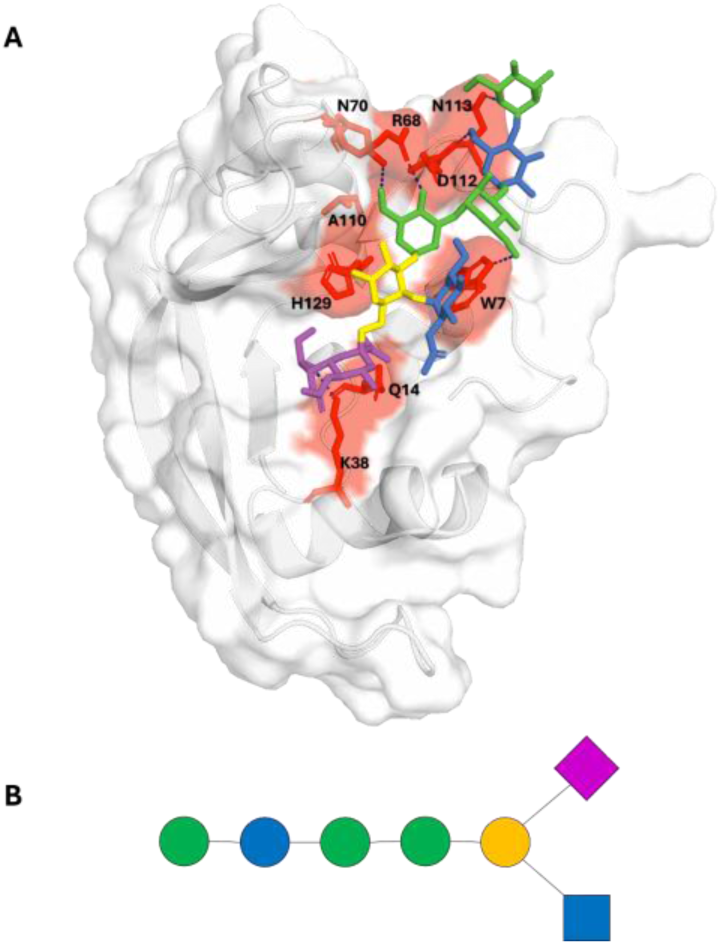
Docking model for EDN-heptasaccharide complex. (A) Position of the heptasaccharide model on the EDN structure. Dashed lines indicate predicted hydrogen bonds. Protein surface and stick representation, where residues interacting with the glycan are shown in red. (B) Schematic representation of the heptasaccharide model following the SNFG convention, in which each monosaccharide is colour-coded to match the corresponding representation in panel A.

For each molecular modelling simulation we selected the model with the lowest affinity. For the MAN-GLC disaccharide at the S_0_–S_1_ region (-3.5 kcal/mol), we observed that this disaccharide fragment preferentially adopted an orientation that overlapped with the GLC position of MAN-GLC observed in the crystal structure. Docking results showed that GLC can hydrogen bond to the S76 side chain, MAN to L45 main chain, and T46, N50, S76, Q77, and V78 can participate in vdW interactions.

For NAG-GLA-SLB docking at the S_5_–S_6_ region, we observed a predominant clustering within two adjacent regions. We prioritized the model with the NAG end close to S_6_, showing the lowest affinity values and the maximum number of Hbond interactions. SLB could form hydrogen bonds with the Ser89 and Cys37 main chains, while NAG preferentially interacted to R35 (Figure S9), with C37, Q91, and N92 potentially participating in vdW interactions.

Finally, by docking the heptasaccharide onto the EDN structure (Figure 8), we obtained result models clustering at the protein nucleotide binding groove, centered around the S_3_ site. After comparing all output models, we selected the model with the second-best predicted affinity energy ranking and H-bond number, which could well cover the S2, S3, and S5 sites, particularly with MAN and SLB positions close to those observed in the protein crystal complexes. We observed that the heptasaccharide SLB unit can H-bond to to K38, MAN to W7, R68, N70, and N113, GLC to D112, while Q14, A110, and H129 can make vdW interactions. Overall, the docked heptamer model is partially reproducing the putative binding of an oligonucleotide spanning from p-_1_ to B_2_ protein sites, shared by both eosinophil RNases (Tables 2 and 3; Figure 7).

## 4. Discussion

It is of particular interest to unravel the sugar binding mode of proteins involved in carbohydrate recognition. A variety of lectins are involved in cell adhesion, communication, signalling, and immune recognition. Unfortunately, both structural determination and prediction of protein complexes with oligosaccharides represent a particular challenge due to glycan diversity, complexity, high molecular sizes and torsion flexibility[43, 44]. In this work we identified 7 sugar binding sites for human RNase 2/EDN. The results support the classification of RNase 2 as a lectin protein and represent the first report of a mammalian lectin within the RNase A family. This is also the first experimental report of RNase 2 sugar binding by MX crystallography. All structural complex data was obtained at atomic resolution allowing us to unambiguously outline the protein surface cavities that specifically accommodate diverse mono and disaccharides. Clustering of all tested molecules in conserved sites highlights the specific residues unique to EDN (Figure 7; Table 3) and provides a valuable information for future structure-functional work towards a better understanding of the protein physiological role.

First, based on a high throughput screening of the protein crystallization conditions, we identified a conserved site for recognition of carboxyl anion groups. We solved 3 crystal structures of RNase 2 in complex with citrate, malonate and tartrate and defined a common binding pocket (A1) (Figure 1). The location of the carboxylic anion molecules overlapped with previous equivalent positioning of acetate and succinate in RNase 2 solved crystal structures (PDB codes:1GQV, 2BZZ and 8F5X) [33, 45]. An equivalent site was also observed in ECP crystal structure in the presence of citrate (PDB code: 4OXF). Moreover, in this study, we observed the presence of acetate at the same location in some crystal structures where this compound was even absent from the crystallization buffer and was only used during the protein purification protocol, emphasizing the strong affinity of both eosinophil RNases (EDN and ECP) towards this carboxylate anion at the described A1 site. Altogether, these data encouraged us to further explore the protein binding to biomolecules that include anion groups with potential biological significance. Next, we hypothesized that the spotted region might be involved in some protein-sugar interaction. Indeed, a quick search by the software *Patchsearch* (https://bioserv.rpbs.univ-paris-diderot.fr/services/PatchSearch/) predicted EDN binding to glucose at a close by location to the identified A1 region.

Trials of protein crystal soaking with a variety of natural sugars proved very successful and we were able to obtain 10 datasets of RNase 2 in complex with mono and disaccharides. Interestingly, several tested monosaccharides (glucose, galactose and fucose) overlapped in a main conserved sugar binding site (S_1_) adjacent to the first spotted A1 anion site (Figures 2 and 3). Proximity of the A1 anion site to the monosaccharide S_1_ site, prompted us to ask whether this region might act as a general N-acetyl–sugar recognition site. We therefore soaked crystals with four N-acetylated sugars of distinct structures (NAG, MUB, NGA and SLB). However, none of these ligands produced additional electron density at the original glucose/acetate site. Instead, they bound at other locations on the protein surface. These observations suggest that, at this region, acetate binding most likely reflects a local affinity for a smaller substituent, whereas full N-acetyl sugars, because of their larger size or different conformational preferences, cannot be accommodated within the same pocket. Besides, prior presence of an acetate bound molecule at the protein A1 site might prevent binding of the added N-acetyl sugar in the crystal soaking solution. Nonetheless, crystal soaking experiments proved successful, and we identified several sugar binding sites, with diverse saccharide type selectivity. Mostly, the tested sugars were bound via polar and vdW contacts, with secondary participation of electrostatic interactions through some specific Arg residues (Table 2).

Another interesting feature is the identification of more than one binding site for some of the tested monosaccharides. For example, we identified four binding sites for the N-acetyl neuraminic acid, indicating that RNase 2 has the potential to anchor sialic acids at multiple positions. Among them, at site S_5_, the negatively charged carboxylate group of SLB forms stabilizing interactions with R35, R68 and R132. Interestingly, these three cationic residues were identified by *ConSurf* analysis as sequence specific for RNase 2 (Figure 7B; Table 3). It is noteworthy that both SLB1 and SLB3 bound at the S_5_ pocket (Figure 5A) are anchored by arginine residues equidistantly spaced, which suggests that the protein might recognize sialylated oligosaccharides, such as chains containing three or more consecutive SLB units.

As commented before, the RNase A superfamily includes a group of frog RNases that were initially described as lectins, before even their enzymatic activity was discovered. A main representant of the frog lectin RNases is cSBL, a Sialic-acid-Binding Lectin isolated from bullfrog (*Rana catesbeiana*) eggs. Here, by EDN crystal soaking with SLB, we identified 4 distinct binding sites (Figure 5). Conserved and non-conserved structural features are likely to underlie differences between cSBL and RNase 2 in sugar specificity and biological properties. *R. catesbeiana* RNase was crystallized in the presence of 4 mM sialic acid[46]. Two bound molecules were identified in cSBL, one close to the active site and the second one bound with lower affinity to a region that would correspond to residues 61-77 in RNase 2, related to the defined S_3_ site in the present work (Table 2). However, this region reveals a significant divergence within the RNase A family [47], with a loop insertion for mammalian RNases, not present in lower order vertebrates. On the other hand, sugar binding in frog RNases was correlated to antitumoral properties. The amphibian proteins were reported to display a pronounced selectivity for tumour cells, where binding to sialylated glycoproteins on the cell surface mediated cell internalization and ultimately induced cell death [48]. In addition, Ogawa and co-workers demonstrated that by increasing the net positive charge by Arg incorporation at the protein surface they enhanced endocytosis and antitumoral activity[49]. Interestingly, the antitumoral properties of frog RNases seemed to be dependent on the sugar type at the cell surface. Indeed, altered patterns of protein glycosylation are widely recognized as an important hallmark of cancer, contributing to tumour growth, metastasis, immune evasion and therapy resistance[50]. On the other hand, the RNases from both *Rana catesbeiana* and *Rana japonica* displayed agglutinating activities for tumour cells but not for erythrocytes, lymphocytes or fibroblasts. The proteins’ cell binding was confirmed to be related to tumour cells sialic acid content, as treatment with sialidase reduced the interaction[9]. Moreover, the binding could be blocked by addition of ganglioside, heparin, spermidine or mucin. Interestingly, pyrimidine nucleotides were also inhibitors of the frog lectin activity [51].

In this context, our observation that EDN can recognize several simple monosaccharides (such as GLA and NAG) as well as multiple SLB sites provides a structural basis for future studies addressing whether the protein preferentially targets glycan features that are common on cancer cells, for example, highly sialylated epitopes, or other glycosylation patterns.

To note, eosinophil RNases were previously reported to bind glycosaminoglycans. Both cationic RNases were observed to bind to heparin when initially purified[17, 52]. Subsequent work on the homologue ECP reported that the protein interacted with Beas-2B cells thorough binding to heparan sulphate, prior to endocytosis and traffic to lysosomal compartment[53]; a process that was mediated by residues 34-38 [23]. A later study identified an equivalent region in EDN (34QRRCKN39) for heparin binding [21]. This region includes a conserved R36 in both ECP and EDN and would correspond to currently identified S_5_ site (Table 3). A second heparin binding domain of EDN was spotted at region 65-70, which corresponds to the present S_3_ sugar binding site. Finally, a third site was reported, at the 113-119 region, which would partially overlap at the S_2_ sugar site.

On the other side, the work by Hung and collaborators using heparin oligosaccharides reported more than 300 times lower calculated IC50 for the inhibition of protein binding to Beas-2B cells when the sugar units increased from three to seven [20]. However, in those earlier studies the main driving force identified relied in the electrostatic interaction between protein cationic groups and sugar sulphate groups. This differs significantly from the present data, where we observe that the studied sugars are mainly bound by polar H-bonds and vdW interactions. Nonetheless, those previous studies provided highly valuable information to infer the optimal length of the saccharide ligand in physiological conditions. Interestingly, both our crystallographic and molecular docking data highlight a partial overlapping between the identified sugar binding sites and the previously described enzyme-substrate binding sites. A good overlapping was also achieved by using the tetranucleotide in complex with RNase A (PDB ID: 1RCN)[54] and a tetrasaccharide modelled into ECP, the latest based on the protein crystal structure in the presence of SO_4_ anions[22]. Accordingly, earlier molecular modelling on ECP binding revealed that the spacing of the protein surface exposed cationic residues could fit both an extended oligomer composed of either sulphated sugars or nucleotides [55], with an equivalent binding mode pattern to interact with either sulphate or phosphate groups respectively. In fact, ECP binding to a heparin trisaccharide was analysed by NMR[19], where the trimer was located at the protein active site, with additional involvement of the 34-39 region (p_-1_ region related to S_5_ sugar site). Moreover, previous experimental data demonstrated that addition of heparin inhibited the enzymatic activity of ECP[18]. In our present work, we also identified some correspondence between EDN sugar and nucleotide binding sites at the protein active site groove, expanding from p_-1_/p_1_/B_1_to B_2_ (S_3_-S_5_) (Table 2). However, apart from the partial overlapping of central S_3_ to S_5_ sites with EDN substrate binding groove, the main sugar binding sites identified in this study (S_0_ to S_6_) are unique and rely not on positively cationic surface regions but on multiple polar and hydrophobic patches (Figure 6, Table 2). Thus, the present study using mono and disaccharides by MX allows a more complete exploration of the protein surface chemical space and offers a robust and unambiguous information on its binding mode to diverse saccharide units.

Evolutionary analysis within the human RNase A family members (Figure 7) reveals that the identified S_1_ is the most characteristic region of EDN and is only partially conserved in EDN/ECP, where several residues are unique (see Table 3). Likewise, we find other sites, as S_0_, S_3_ and S_6_, which also present unique traits for EDN, together with S_4_ and S_5_ with shared EDN/ECP structural determinants. Analysis of EDN conserved-variable residues highlights some unique surface exposed regions (Figure 7B). Among them, R132 binding to SLB at S_5_ (Table 3) is an EDN-specific residue that was previously identified as a key determinant to distinguish EDN from ECP[56]. Whereas H82, which is shared by EDN and ECP, makes vdW contacts with SLB at S_5_. We also find the unique R68, which hydrogen-bonds with both MAN and TRE at the S_3_ pocket. Last, we would like to highlight L45, which is unique to EDN sequence (Figure 7A) and participates in S_0_ and S_1_, mostly recognizing mannose, glucose, galactose or fucose. Another residue of interest identified in the evolutionary analysis of eosinophil RNases when exploring their divergence from a common ancestor, is R132, which was correlated to the increased antiviral activity of EDN [45, 56]. Interestingly, the recently published crystal structure of *Macaca fascicularis* ECP in the presence of citrate highlights essential differences at the A1-S_1_ environment (PDB ID: 7TY1) [45]. In particular, we observe conserved interactions at Y107 but a reduced hydrophobic patch at the 45-46 region. On the other hand, the S_3_ site is located at the L4-L7 loop interface. According to Doucet and colleagues molecular dynamics studies, L4 (61-70aa) is the main flexible loop in EDN and its proximity to L7 (114-124aa), a loop characteristic of eosinophil RNases, can determine the protein functionality [16, 45]. Comparative sugar binding within the RNase A family was also explored by Raines and coworkers using heparin and RNase1 as a model. They studied representative RNase1 members and reported an increase in heparin binding in higher versus lower order vertebrates [57]. We could speculate that the eosinophil RNases branch, which has been the last to evolve within the family (Figure 7C), could have acquired an enhanced GAG binding affinity and evolved towards a better recognition and adherence to hominoid immune cell surface. Therefore, characterization of EDN selective sugar recognition pattern is of particular interest to understand the biology of the hominoid primate branch of RNases.

To note, neither the previous reported frog leczymes nor the current sugar-binding pattern identified in EDN conform to any of the canonical carbohydrate-recognition domain (CRD) motifs described in lectins. Although there is no universal consensus for carbohydrate recognition, most common lectins employ divalent metal ions (Ca²⁺, Mg²⁺) to mediate interactions with sugar-ring hydroxyls. In our case, no metal ions are involved, and the RNase 2 sites do not fall into the classical C-type CRD category[58].

Last, the recent discovery of glycosylated RNAs, exposed at the blood cell type surfaces[25, 59], provide an additional layer of complexity in cell immune modulation. We can speculate that glycosyl binding sites adjacent to the RNase active site may help the protein to target specifically the GlycoRNAs exposed at the cell surface. Alternatively, RNA anchored at the cell membrane might have acquired glycosylation to protect themselves from the presence of circulating extracellular endonucleases[60]. GlycoRNAs differ according to blood cell type and are involved in specific cell-cell interactions, so disruption of these connecting molecules might have key implications in the immune system[61].

In summary, identifying the protein’s selective sugar binding mode is of utmost importance due to its crucial role on surface glycan composition in cell-cell recognition, host-pathogen interactions or discrimination between healthy and tumoral cells.

In this work we were able to define 7 sugar binding pockets at EDN protein surface. Further work is planned to complement the present structural work with some biophysical characterization and functional assays. Throughput screening experiments will also be needed to find out the best sugar partners for EDN in physiological conditions. Identification of EDN sweet partners *in vivo* is definitely of great interest for the development of novel therapies.

## DATA AVAILABILITY

Coordinates and structure factors have been deposited at the *PDB* server under the accession codes 8QEW, 9GNT, 9GO7, 9GO8, 9GO4, 9GQ0, 9GQM, 9GQP, 9QYO, 9QR7, 9R6D, 9RAN and 9R6E.

## ACKNOWLEDGEMENTS

We gratefully acknowledge on-site support from Dr. Roeland Boer, Dr. Isidro Crespo, Dr. Fernando Gil and Dr. Aleix Tarrés Solé at beamline BL13 ALBA synchrotron. XK and JL were recipients of a predoctoral *China Scholar Fellowship*. GP-E was a recipient of a *Margarita Salas* postdoctoral fellowship. Research was support by *Agencia Estatal de Investigación* (PID2022-137872NB-I00).

## AUTHOR CONTRIBUTION

XK, JL and EB: Conceptualization, Formal analysis, Methodology, Validation, Writing—original draft, review and editing. GPE: Formal analysis, Methodology, original draft review.

## SUPPLEMENTARY DATA

**Figure S1.** Electron density and interaction analysis of carboxylate anions bound to EDN.

**Figure S2.** Second anion binding site (A2) identified in EDN complexes.

**Figure S3.** Binding sites of tested monosaccharides and disaccharides on the surface of EDN.

**Figure S4.** Fucose and galactose recognition at the EDN S_1_ sugar-binding site.

**Figure S5.** Location of EDN sugar binding sites S0 and S4.

**Figure S6.** Sugar-binding site S_2_ for Mannose (MAN), N-acetyl-beta-neuraminic acid (SLB) and N-acetylmuramic acid (MUB).

**Figure S7.** Binding interaction of N-acetylated sugar derivatives on the surface of EDN.

**Figure S8.** Disaccharide MAN-GLC docked into the S1 pocket.

**Figure S9.** Binding sites of trisaccharide NAG-GLA-SLB docked into S5 and S6 pockets.

**Table S1.** Optimization ranges tested for the three crystallization conditions selected from the initial screening hits.

**Table S2.** Crystallization conditions and X-ray data collection/refinement statistics for EDN structure complexes.

**Table S3.** Grid box parameters used for molecular docking simulations.

**Table S4.** Estimated binding energies for docking of mono and disaccharides into EDN sugar-binding pockets using *Autodock vina-carb*.

