## Supplemental figures and tables for "Structural basis for saccharide binding by human RNase 2/EDN, a protein combining enzymatic and lectin properties"

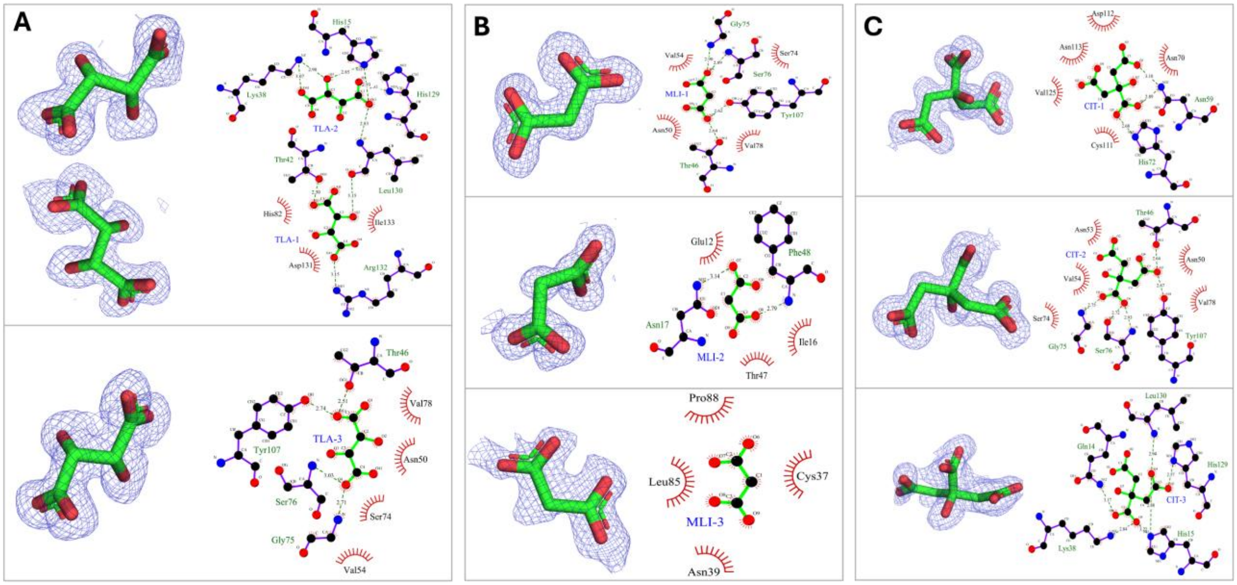


Tartrate Malonate Citrate

**Figure S1**. **Electron density and interaction analysis of carboxylate anions bound to EDN**. The panels highlight the anion-binding sites, displaying 2FoFc electron density maps for the ligands and detailing their interactions with protein residues, as depicted by LigPlot. (A) Tartrate (TLA) ; (B) Malonate (MLI) and (C) Citrate (CIT).

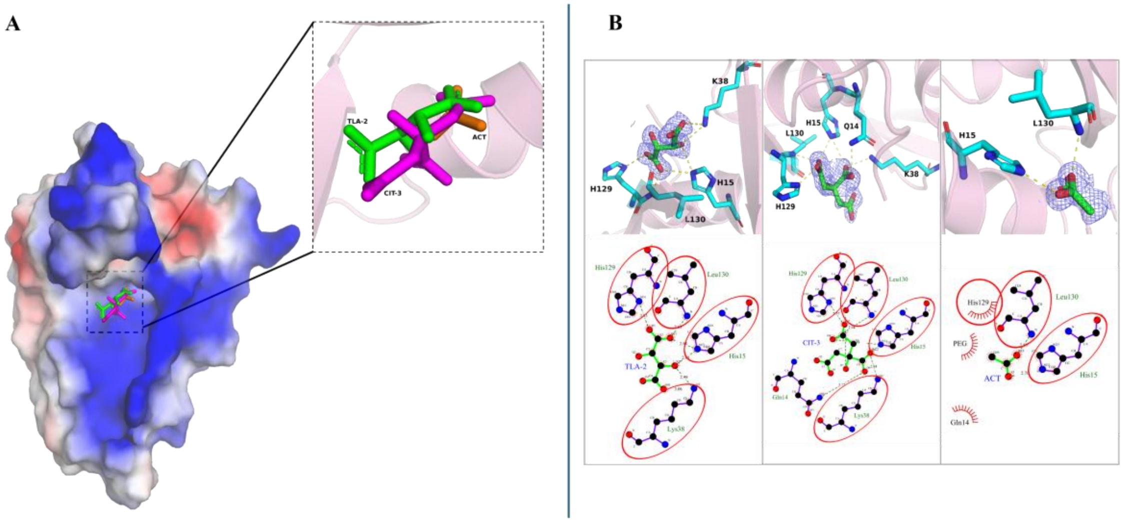


**Figure S2. Second anion binding site (A2) identified in EDN complexes**. (A) Overall view showing the ligand clustering at the A2 site. (B) Close-up view of anion binding at the $S_2$ site and Ligplot interaction diagrams. Interacting residues are displayed as sticks, Residues highlighted by red ellipses are consistently observed across different ligand complexes, identifying them as common binding hotspots.

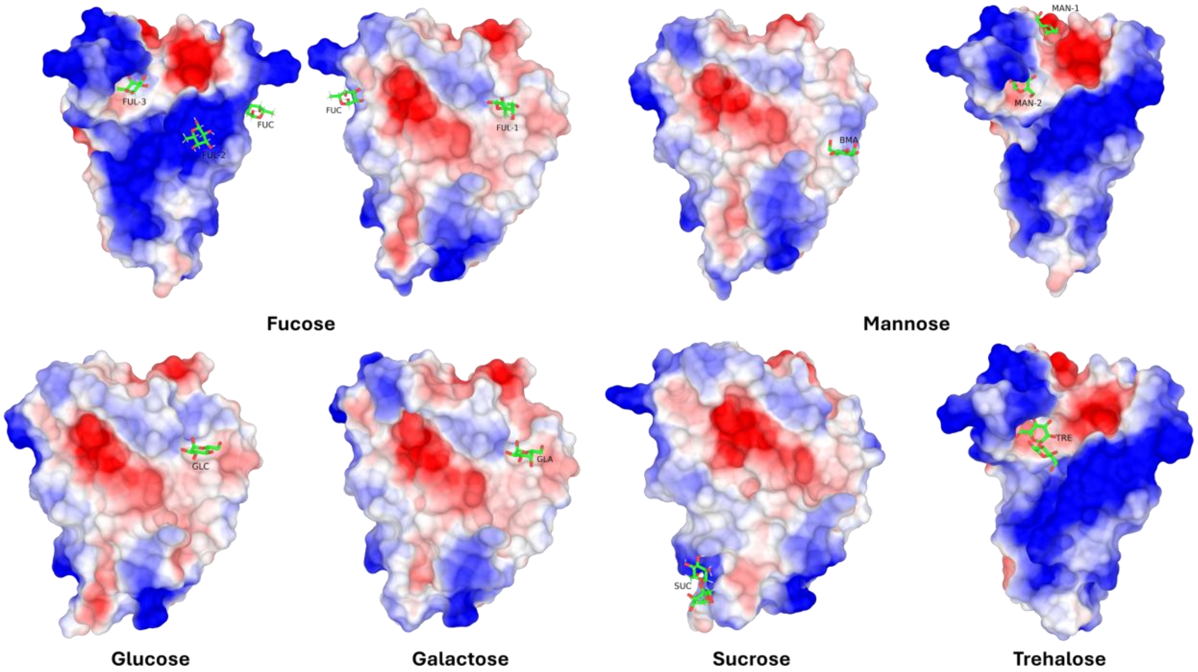


**Figure S3. Binding sites of tested monosaccharides and disaccharides on the surface of EDN.** The molecular surface of the EDN protein is represented by the electrostatic potential to illustrate the charge distribution across the binding pockets. Positively charged regions are shown in blue, negatively charged regions in red, and neutral regions in white. The carbohydrate ligands are depicted as stick models.

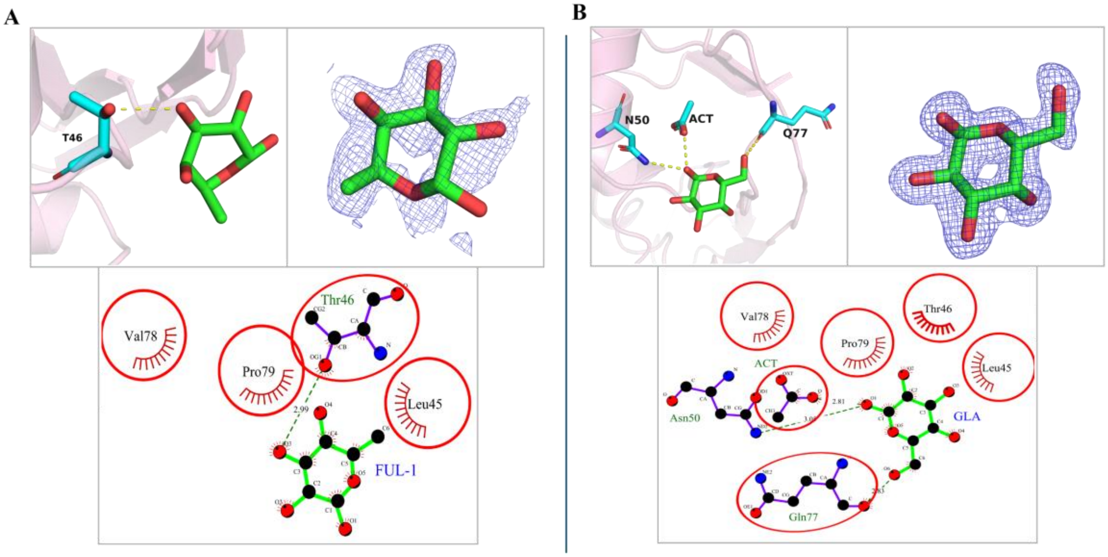


**Figure S4**. **Fucose and galactose recognition at the EDN $S_1$ sugar-binding site.** (A) Close-up view and electron density map of fucose (FUL-1) bound at $S_1$, (B) Close-up view and interaction diagram of galactose (GLA) at $S_1$.

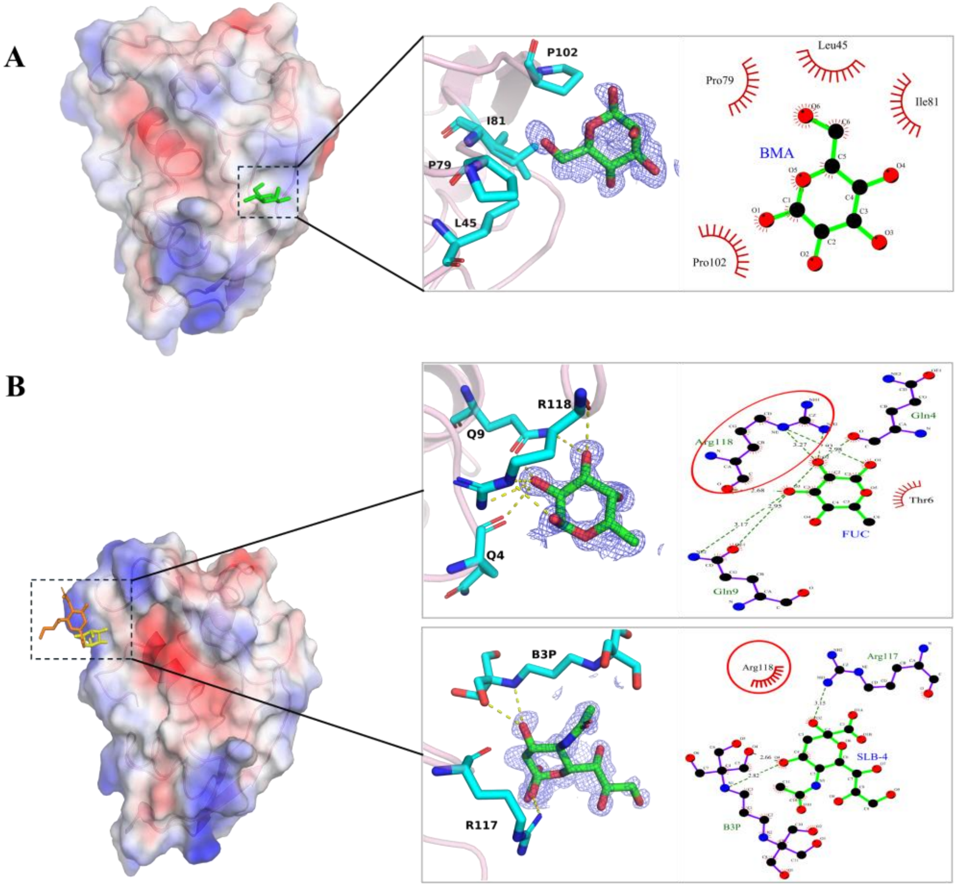


**Figure S5. Location of EDN sugar binding sites $S_0$ and $S_4$.** (A) Location of $S_0$ site on EDN surface. (B) Location of $S_4$ site on EDN surface. The $S_0$ site includes mannose BMA, and the $S_4$ site includes N-acetyl-beta-neuraminic acid (SLB4) and fucose (FUC). The dashed lines represent hydrogen bonds, and the red circles indicate amino acids common to these binding sites. Red arcs representing hydrophobic interactions.

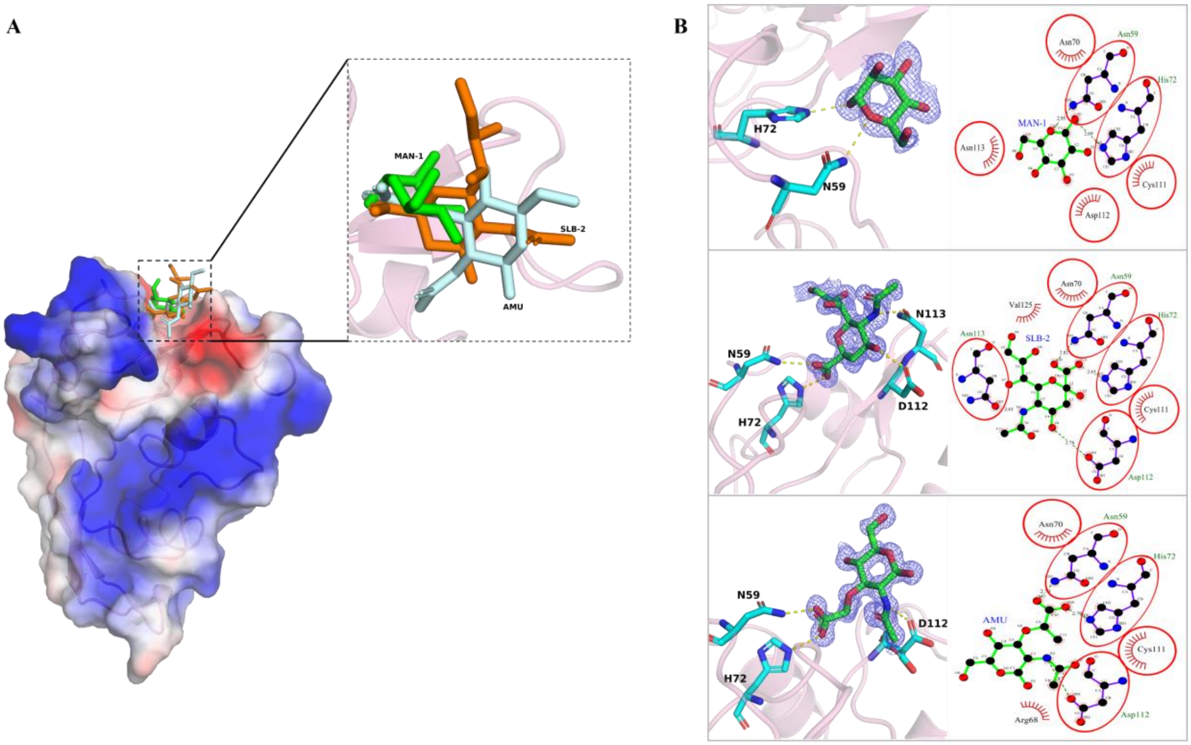


**Figure S6. Sugar-binding site $S_2$ for Mannose (MAN), N-acetyl-beta-neuraminic acid (SLB) and N-acetylmuramic acid (AMU).** (A) Overall view showing the clustering of the three sugars at the $S_2$ site. (B) Close-up view of sugar binding at the S2 site and Ligplot interaction diagrams. Interacting residues are displayed as sticks; residues highlighted by red ellipses are consistently observed across different ligand complexes, identifying them as common binding hotspots.

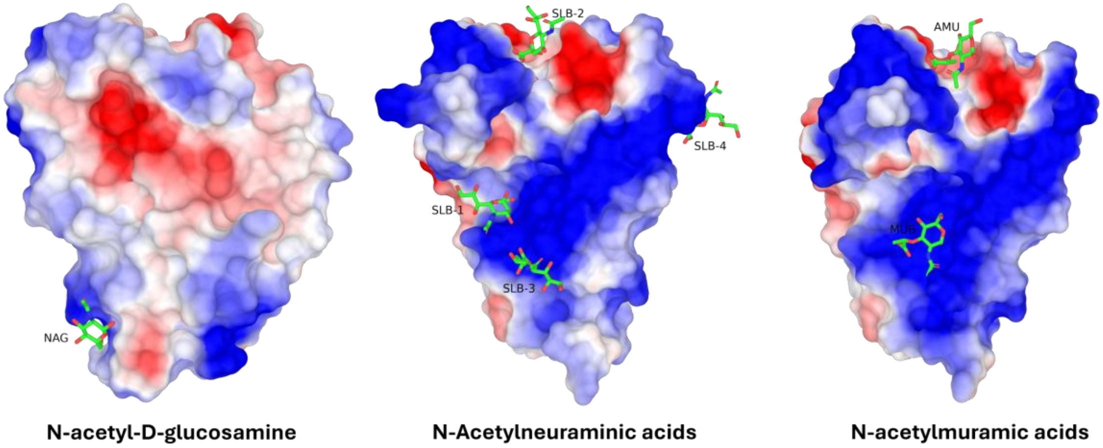


**Figure S7. Binding interaction of N-acetylated sugar derivatives on the surface of EDN.** The EDN protein surface is represented by its electrostatic potential to visualize the charge complementarity at the binding interface (blue: positive; red: negative; white: neutral). The specific ligands shown, rendered as stick models, are, N-acetyl-D-glucosamine (NAG), N-acetylneuraminic acid (SLB 1 to 4) and N-α and β acetylmuramic acid (MUB and AMU).

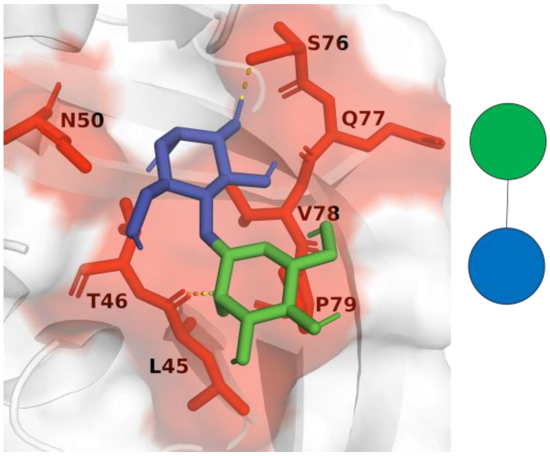


**Figure S8**. **Predicted location of MAN-GLC disaccharide docked into the $S_1$ pocket.** Mannose and glucose are indicated following the SNFG (Symbol Nomenclature for Glycans) convention, where the green circle represents mannose and the blue circle represents glucose. The protein surface coloured in red highlights the amino acids participating in both hydrogen bond and vdW interactions.

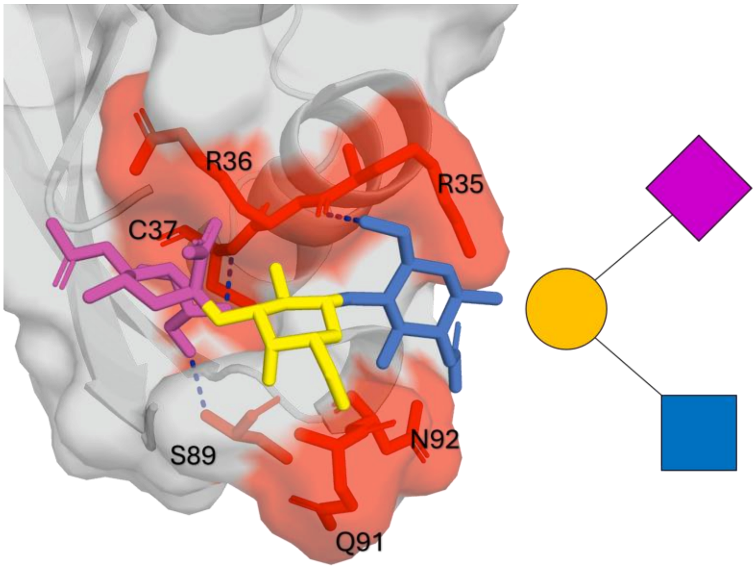


**Figure S9**. **Predicted binding of NAG-GLA-SLB trisaccharide docked into $S_5$ and $S_6$ pockets.** NAG-GLA-SLB are indicated on the right following the SNFG (Symbol Nomenclature for Glycans) convention, where N-acetyl-beta-neuraminic acid (SLB) is coloured in purple; Galactosamine (GLA) is coloured in yellow; and N-Acetyl-D-glucosamine (NAG) in blue. The protein surface coloured in red highlights the amino acids participating in both hydrogen bond and vdW interactions.

**Table S1. Optimization ranges tested for the three crystallization conditions selected from the initial screening hits.** The table summarizes the pH values and concentration of precipitants explored during the refinement of conditions 1–3 in order to obtain reproducible crystals suitable for ligand soaking and X-ray diffraction.

| Condition | Variable Components | Tested Range | Fixed Components |
|---|---|---|---|
| **Condition 1**<br>0.2 M Sodium formate<br>0.1 M Bis-Tris propane pH 8.5<br>20% PEG 3350<br>(From H6 of the PACT plate) | PEG 3350 | 14–24% (w/v)<br>(14, 16, 18, 20, 22, 24%) | Sodium formate 0.2 M<br>Bis-Tris propane 0.1 M<br>pH 8.5 |
| | Sodium formate | 0.15–0.30 M<br>(0.15, 0.18, 0.20, 0.22, 0.25, 0.30 M) | PEG 3350 20%<br>Bis-Tris propane 0.1 M<br>pH 8.5 |
| | pH | 7.8–9.0<br>(7.8, 8.0, 8.2, 8.5, 8.7, 9.0) | Sodium formate 0.2 M<br>Bis-Tris propane 0.1 M<br>PEG 3350 20% |
| | Combined variations | PEG + pH<br>PEG + Salt<br>Salt + pH | — |
| **Condition 2**<br>0.1 M Sodium acetate pH 7.0<br>12% PEG 3350<br>(From G1 of the PEG HR2-139 plate) | PEG 3350 | 8–16% (w/v)<br>(8, 10, 12, 14, 15, 16%) | Sodium acetate 0.1 M<br>pH 7 |
| | pH | 6.5–7.8<br>(6.5, 6.8, 7.0, 7.2, 7.5, 7.8) | PEG 3350<br>Sodium acetate 0.1 M |
| | Combined variations | PEG3350(8,10,14)<br>+pH(6.5, 6.8, 7.2, 7.5) | — |
| **Condition 3**<br>0.1 M HEPES pH 7.5<br>10% PEG 6000<br>5% MPD<br>(From E12 of the Structure plate) | PEG 6000 | 8–15% (w/v)<br>(8, 9, 10, 12, 14, 15%) | 0.1M HEPES buffer<br>5% MPD<br>pH 7.5 |
| | MPD | 3–10% (v/v)<br>(3, 4, 5, 7, 9, 10%) | 0.1M HEPES buffer<br>10% PEG 6000<br>pH 7.5 |
| | pH | 7.0–8.2<br>(7.0, 7.2, 7.5, 7.7, 8.0, 8.2) | 0.1M HEPES buffer<br>10% PEG 6000<br>5% MPD |
| | Combined variations | PEG 6000(8, 10, 15)<br>+pH (7.2, 7.5, 7.7)<br>+MPD(3, 5, 10%) | 0.1M HEPES buffer |

**Table S2.** Crystallization conditions and X-ray data collection/refinement statistics for EDN structure complexes.

| Dataset name | CIT | MLI | TLA | GLC+CIT | SUC | FUC | GLC+FUC |
|---|---|---|---|---|---|---|---|
| **PDB ID** | **9GO4** | **9GO8** | **8QEW** | **9GNT** | **9GO7** | **9GQ0** | **9GQM** |
| **Resolution (Å)** | 38.41-1.18 (1.26-1.18) | 33.59-1.04 (1.08-1.04) | 38.5-1.197 (1.25-1.2) | 33.71-1.31 (1.39-1.31) | 38.51-1.04 (1.1-1.04) | 38.48-1.04 (1.08-1.04) | 38.60-1.01 (1.07-1.01) |
| **a, b, c (Å)** | 41.7 52.4 56.5 | 41.8 52.7 56.5 | 41.8 52.7 56.4 | 41.9 53.0 56.8 | 41.8 52.6 56.6 | 41.7 52.6 56.5 | 41.9 52.8 56.5 |
| **α, β, γ (o)** | 90 90 90 | 90 90 90 | 90 90 90 | 90 90 90 | 90 90 90 | 90 90 90 | 90 90 90 |
| **Total reflections** | 130685(2166) | 301182(5701) | 176318(2629) | 111427(6734) | 188571(6687) | 374858(6298) | 305409(4705) |
| **Unique reflections** | 32960 (1636) | 53346 (2667) | 34211 (1711) | 25888 (1296) | 46578 (2330) | 55066 (2753) | 56650 (2832) |
| **Data redundancy** | 4 | 5.6 | 5.2 | 4.3 | 4 | 6.8 | 5.4 |
| **Multiplicity** | 4.0(1.3) | 5.6(2.1) | 5.2(1.5) | 4.3(5.2) | 4.0(2.9) | 6.8(2.3) | 5.4(1.7) |
| **Completeness (%)** | 88.5 (42.8) | 90.3 (45.2) | 89.4(42.1) | 90.9 (46.5) | 83.6 (30.6) | 91.8(47.2) | 86.9 (30.4) |
| **Mean I/s(I)** | 11.7 (2) | 21.3 (2.7) | 13.8(1.90) | 8.1 (1.7) | 23.9 (5.5) | 22.6 (2) | 18.5 (3.3) |
| **Wilson B-factor** | 9.85 | 7.16 | 8.5 | 10.27 | 8.52 | 10.9 | 7.86 |
| **Rmerge** | 0.069(0.348) | 0.054(0.345) | 0.066(0.341) | 0.170(1.354) | 0.031(0.147) | 0.038(0.361) | 0.051(0.161) |
| **CC1/2** | 0.998(0.794) | 0.999(0.756) | 0.998(0.825) | 0.994(0.425) | 0.999(0.967) | 0.999(0.787) | 0.998(0.932) |
| **Rwork** | 0.1516 (0.2585) | 0.1361 (0.2783) | 0.1770 (0.2892) | 0.1613 (0.2837) | 0.1226 (0.1374) | 0.1305 (0.2432) | 0.1351 (0.1761) |
| **Rfree** | 0.1879 (0.3384) | 0.1641 (0.3181) | 0.1770 (0.2892) | 0.2147 (0.3677) | 0.1412 (0.1950) | 0.1416 (0.22650 | 0.1506 (0.2089) |
| **Nº of non-H atoms** | 1425 | 1454 | 1280 | 1388 | 1440 | 1372 | 1426 |
| **Protein** | 1156 | 1186 | 1098 | 1125 | 1175 | 1121 | 1171 |
| **Ligands** | 59 | 56 | 30 | 53 | 62 | 61 | 48 |
| **Solvent** | 210 | 233 | 152 | 210 | 203 | 199 | 220 |
| **Prot. residues** | 135 | 136 | 135 | 135 | 135 | 135 | 135 |
| **RMS (bonds)** | 0.006 | 0.008 | 0.006 | 0.006 | 0.006 | 0.005 | 0.005 |
| **RMS (angles)** | 0.94 | 0.99 | 0.95 | 1 | 0.98 | 0.88 | 1.18 |

| | | | | | | | |
|---|---|---|---|---|---|---|---|
| **Ramachandran favored (%)** | 98.5 | 98.5 | 98.5 | 99.25 | 97.74 | 100 | 100 |
| **Ramachandran allowed (%)** | 1.5 | 1.5 | 1.5 | 0.75 | 2.26 | 0 | 0 |
| **Ramachandran outliers (%)** | 0 | 0 | 0 | 0 | 0 | 0 | 0 |
| **Rotamer outliers (%)** | 0.74 | 0 | 0.78 | 0 | 0.72 | 0.77 | 0 |
| **Clashscore** | 1.69 | 6.7 | 0 | 4.37 | 0.82 | 0.87 | 2.94 |
| **Average B-factor** | 17.76 | 13.43 | 12.43 | 18.68 | 14.49 | 19.83 | 14.18 |
| **macromolecules** | 13.98 | 10.25 | 11.52 | 14.04 | 10.72 | 14.17 | 10.77 |
| **Ligands** | 24.5 | 16.68 | 14.74 | 31.41 | 25.42 | 38.69 | 17.35 |
| **Solvent** | 36.6 | 29.13 | 18.58 | 40.33 | 32.97 | 45.67 | 31.85 |
| **Crystallization buffer** | 0.2 M Ammonium citrate dibasic, 20% PEG 3350, pH 5.1 | 0.2 M Sodium malonate,20% PEG 3350, pH 4 | 0.2 M Ammonium tartrate dibasic,20% PEG3350, pH 7 | 0.05 M Citric acid, 0.05 M BIS-TRIS propane, 16% Polyethylene glycol 3350, pH 5 | 0.2M Sodium format, 0.1M Bis-Tris, 18% PEG3350, pH 8.2 | 0.18M Sodium Format, 0.1M Bis-Tris propane, 15% PEG 3350, pH 8.2 | 0.1M sodium acetate, 10% PEG3350, pH 7.2 |
| **Soaking Ligand Conc.** | - | - | - | 250mg/ml | 250mg/ml | 250mg/ml | 250mg/ml |
| **Soaking Time** | - | - | - | 30min | 1h | 1h | 30min |

| **Protein** | **SLB** | **MUB** | **NAG** | **GLA** | **MAN** | **TRE** |
|---|---|---|---|---|---|---|
| **PDB ID** | **9GQP** | **9QR7** | **9R6E** | **9RAN** | **9QYO** | **9R6D** |
| **Resolution (Å)** | 38.67-1.01 (1.08-1.01) | 38.58-1.02 (1.07-1.02) | 38.51-1.02 (1.09-1.02) | 38.51-1.02 (1.07-1.02) | 33.68-1.02 (1.1-1.02) | 38.56-0.94 (0.99-0.94) |
| **a, b, c (Å)** | 41.9 53.1 56.5 | 41.7 52.9 56.4 | 41.8 52.6 56.5 | 41.7 52.7 56.5 | 41.8 52.6 56.9 | 42.0 52.5 56.8 |
| **α, β, γ (o)** | 90 90 90 | 90 90 90 | 90 90 90 | 90 90 90 | 90 90 90 | 90 90 90 |
| **Total reflections** | 308631(5659) | 254783(4282) | 213887(5017) | 215437(3797) | 203285(5851) | 398018(5932) |
| **Unique reflections** | 54023(2701) | 55704(2785) | 49527(2476) | 54608(2730) | 49102(2455) | 72603(3630) |
| **Data redundancy** | 5.7 | 4.6 | 4.3 | 3.9 | 4.1 | 5.5 |
| **Multiplicity** | 5.7(2.1) | 4.6(1.5) | 4.3(2) | 3.9(1.4) | 4.1(2.4) | 5.5(1.6) |
| **Completeness (%)** | 85.5(32.6) | 88.4(34.6) | 83(30.8) | 84.3(27.6) | 78.4(22.5) | 90.3(36.5) |
| **Mean I/s(I)** | 21(2.9) | 16.8(2.5) | 15.5(3) | 29.6(7.8) | 20.8(10) | 19.9(2.3) |
| **Wilson B-factor** | 7.96 | 8.64 | 8.08 | 8.06 | 8.12 | 8.71 |
| **Rmerge** | 0.044(0.197) | 0.045(0.221) | 0.05(0.397) | 0.033(0.056) | 0.063(0.094) | 0.062(0.295) |
| **CC1/2** | 0.998(0.922) | 0.999(0.894) | 0.998(0.709) | 0.995(0.991) | 0.993(0.980) | 0.995(0.835) |
| **Rwork** | 0.1361 (0.1894) | 0.1330 (0.2038) | 0.1264(0.3118) | 0.1390 (0.1366) | 0.1346 (0.1310) | 0.1390 (0.2248) |
| **Rfree** | 0.1504 (0.2091) | 0.1495 (0.1809) | 0.1369(0.3189) | 0.1516 (0.1211) | 0.1496 (0.1902) | 0.1477 (0.3244) |
| **Nº of non-H atoms** | 1494 | 1460 | 1384 | 1427 | 1446 | 1433 |
| **Protein** | 1133 | 1186 | 1124 | 1173 | 1192 | 1196 |
| **Ligands** | 107 | 58 | 60 | 56 | 56 | 38 |
| **Solvent** | 254 | 216 | 200 | 198 | 198 | 199 |
| **Prot. residues** | 135 | 135 | 135 | 135 | 135 | 135 |
| **RMS (bonds)** | 0.006 | 0.005 | 0.006 | 0.005 | 0.005 | 0.005 |
| **RMS (angles)** | 0.89 | 0.89 | 0.94 | 0.83 | 0.82 | 0.86 |
| **Ramachandran favored (%)** | 99.25 | 98.5 | 97.74 | 100 | 97.74 | 99.25 |
| **Ramachandran allowed (%)** | 0.75 | 1.5 | 2.26 | 0 | 2.26 | 0.75 |

| | | | | | | |
|---|---|---|---|---|---|---|
| **Ramachandran outliers (%)** | 0 | 0 | 0 | 0 | 0 | 0 |
| **Rotamer outliers (%)** | 0 | 0.72 | 0 | 0.7 | 0 | 0.75 |
| **Clashscore** | 0.82 | 1.6 | 2.55 | 1.23 | 0.41 | 0.82 |
| **Average B-factor** | 17.5 | 16.19 | 16.09 | 16.84 | 16.23 | 15.43 |
| **macromolecules** | 10..65 | 11.2 | 10.54 | 11.67 | 11.49 | 11.41 |
| **Ligands** | 26.5 | 30.04 | 31.46 | 20.13 | 29.6 | 35.56 |
| **Solvent** | 44.26 | 39.9 | 42.63 | 46.53 | 40.97 | 35.77 |
| **Crystallization buffer** | 0.22M Sodium formate, 0.1M Bis-Tris propane, 12% PEG 3350, pH 7 | | | 0.18M Sodium Formate, 0.1M Bis-Tris propane, 15% PEG 3350, pH 8.2 | 0.22M Formate, 0.1M Bis-Tris propane, 22% PEG 3350, pH 8.5 | 0.1M Sodium formate, 14% PEG 3350, pH 7.2 |
| **Soaking Ligand Conc.** | 50mg/ml | | | 250mg/ml | 250mg/ml | 250mg/ml |
| **Soaking Time** | 1-2 day | | | 12h | 12h | 30min |

**Table S3. Grid box parameters used for molecular docking simulations**. The table lists the center coordinates and box dimensions defining the docking search space for each ligand. All values are reported in Ångström (Å) and correspond to the parameters used in the AutoDock Vina-Carb configuration files.

| Common Name | Ligand | Grid Box Center (Å) | | | Grid Box Size (Å) | | |
|---|---|---|---|---|---|---|---|
| | | x | y | z | x | y | z |
| Fucose | FUC | -3.21 | 14.30 | 11.26 | 13.13 | 13.13 | 13.13 |
| | FUL-1 | 13.95 | -0.00 | 4.05 | 10.60 | 10.60 | 10.60 |
| | FUL-2 | 1.57 | 15.74 | 15.29 | 14.25 | 14.25 | 14.25 |
| | FUL-3 | 12.57 | 7.18 | 22.26 | 14.25 | 14.25 | 14.25 |
| Galactose | GLA | 15.45 | -2.03 | 5.34 | 20.17 | 13.45 | 13.45 |
| Glucose | GLC | 17.13 | -0.89 | 5.34 | 18.17 | 12.11 | 12.11 |
| Mannose | MAN-1 | 5.43 | 0.48 | 23.10 | 17.08 | 15.38 | 15.03 |
| | MAN-2 | 11.58 | 7.88 | 22.98 | 13.67 | 13.67 | 13.67 |
| | BMA | 18.98 | 4.19 | -0.11 | 16.53 | 14.88 | 14.54 |
| Sucrose | SUC | 11.39 | 26.07 | 3.60 | 16.35 | 19.08 | 17.56 |
| Trehalose | TRE | 9.96 | 8.28 | 24.30 | 21.80 | 19.08 | 23.62 |
| N-acetyl-D-glucosamine | NAG | 11.39 | 26.07 | 3.60 | 16.35 | 19.08 | 17.56 |
| N-Acetylmuramic acid | MUB | 14.98 | 21.57 | 10.33 | 19.05 | 19.40 | 18.34 |
| | AMU | 6.14 | 2.00 | 24.41 | 17.64 | 16.58 | 16.23 |
| N-acetylneuraminic acid | SLB-1 | 16.37 | 13.52 | 12.47 | 18.17 | 19.08 | 19.68 |
| | SLB-2 | 5.35 | 0.09 | 22.11 | 18.17 | 19.08 | 19.68 |
| | SLB-3 | 14.69 | 22.63 | 8.50 | 16.83 | 17.68 | 15.43 |
| | SLB-4 | -9.09 | 9.80 | 14.49 | 14.27 | 17.24 | 16.05 |

**Table S4.** Estimated binding energies for docking of mono and disaccharides into EDN sugar-binding pockets using *Autodock vina-carb*.

| **Sugar** | **Estimated affinity** (kcal/mol) | | | |
|---|---|---|---|---|
| FUC | FUC(-3.3) | FUL-1(-3.1) | FUL-2(-3.3) | FUL-3(4.7) |
| GLA | -4.4 | | | |
| GLC | -3.4 | | | |
| MAN | MAN-1(-4.4) | MAN-2(-4.5) | BMA(-3.3) | |
| SUC | -4.1 | | | |
| TRE | -5.2 | | | |
| SLB | SLB-1(-5.8) | SLB-2(-5) | SLB-3(-4.4) | SLB-4(-3.7) |
| NAG | -4.2 | | | |
| MUB | MUB(-5.4) | AMU(-5) | | |

For each sugar, the numbering of docked molecules (e.g. FUL-1–FUL-3, SLB-1–SLB-4) corresponds to the individual sugar molecules identified in the solved crystal structures. The value in parentheses indicates the highest affinity estimated at that site by molecular docking.
